# Molecular determinants of antagonist interactions with chemokine receptors CCR2 and CCR5

**DOI:** 10.1101/2023.11.15.567150

**Authors:** John R.D. Dawson, Grant M. Wadman, Penglie Zhang, Andrew Tebben, Percy H. Carter, Siyi Gu, Thomas Shroka, Leire Borrega-Roman, Catherina L. Salanga, Tracy M. Handel, Irina Kufareva

## Abstract

By driving monocyte chemotaxis, the chemokine receptor CCR2 shapes inflammatory responses and the formation of tumor microenvironments. This makes it a promising target in inflammation and immuno-oncology; however, despite extensive efforts, there are no FDA-approved CCR2-targeting therapeutics. Cited challenges include the redundancy of the chemokine system, suboptimal properties of compound candidates, and species differences that confound the translation of results from animals to humans. Structure-based drug design can rationalize and accelerate the discovery and optimization of CCR2 antagonists to address these challenges. The prerequisites for such efforts include an atomic-level understanding of the molecular determinants of action of existing antagonists.

In this study, using molecular docking and artificial-intelligence-powered compound library screening, we uncover the structural principles of small molecule antagonism and selectivity towards CCR2 and its sister receptor CCR5. CCR2 orthosteric inhibitors are shown to universally occupy an inactive-state-specific tunnel between receptor helices 1 and 7; we also discover an unexpected role for an extra-helical groove accessible through this tunnel, suggesting its potential as a new targetable interface for CCR2 and CCR5 modulation. By contrast, only shape complementarity and limited helix 8 hydrogen bonding govern the binding of various chemotypes of allosteric antagonists. CCR2 residues S101^2.63^ and V244^6.36^ are implicated as determinants of CCR2/CCR5 and human/mouse orthosteric and allosteric antagonist selectivity, respectively, and the role of S101^2.63^ is corroborated through experimental gain-of-function mutagenesis. We establish a critical role of induced fit in antagonist recognition, reveal strong chemotype selectivity of existing structures, and demonstrate the high predictive potential of a new deep-learning-based compound scoring function. Finally, this study expands the available CCR2 structural landscape with computationally generated chemotype-specific models well-suited for structure-based antagonist design.

## Introduction

The human immune system is a network of specialized cells intricately coordinated to protect the body against injury and infection, yet its dysregulation causes inflammatory and autoimmune disorders. Transmembrane chemokine receptors orchestrate the migration of a wide variety of immune cells to sites of injury and inflammation via interactions with their endogenous ligands, the chemokines [1]. Due to their involvement in cancer, inflammation, and other disorders, these Class A G protein-coupled receptors (GPCRs) are considered promising drug targets [2–4]. Notably, the CC chemokine receptor 2 (CCR2), together with its chemokine agonist CCL2, demonstrated potential as a target in cardiovascular disease [5], diabetes [6–9], liver fibrosis [10,11], neuropathic pain [12], rheumatoid arthritis [13], neuroinflammation [14,15], and carcinogenesis [16–18]. In the context of immuno-oncology, CCR2 inhibition has been investigated as a strategy for alleviating immunosuppression and improving the efficacy of chemo-, radiation, and immunotherapies [19–25].

Motivated by these disease associations, industry- and academia-based drug discovery efforts over the last few decades led to the discovery of thousands of small molecule inhibitors of CCR2 and its closest human homologue, CCR5 [26–29]. These compounds are highly diverse from the standpoints of structural features, mechanisms of receptor inhibition [30–32], and pharmacological properties. However, small molecules and biologics targeting the CCR2-CCL2 axis have shown mixed results in early stage clinical trials [6,9,33–42] and none have reached regulatory approval, whereas for CCR5, the only regulatory approval to date has been **maraviroc**, for the treatment of HIV infection [43–46].

Suboptimal properties of drug candidates have been cited as major contributors to clinical failures [29]. At least in inflammatory contexts, therapeutic efficacy of chemokine receptor inhibitors has been linked to high *in vivo* receptor occupancy over extended periods of time [47,48]. This is a challenging task, given that CCR2 antagonists have to competitively inhibit the high-affinity (sub-nanomolar) protein-protein interaction between the receptor and the abundant chemokine. To further complicate the situation, all known inhibitors of the CCR2-CCL2 system result in an apparent “CCL2 induction” where plasma chemokine increases far above pre-treatment levels. Although this phenomenon has been known for a long time [33–35,39], only recently have we started to understand its convoluted biological mechanisms which involve constitutive production of CCL2 by tissues, constitutive uptake and intracellular degradation (“scavenging”) of the chemokine by CCR2, and the inhibition of such scavenging by drugs [49,50]. Importantly, when monitored in mice over time, the CCL2 peak in the blood occurs **after** the peak plasma concentration of the inhibitor [49]. This may have a profound impact on the efficacy of CCR2-targeted therapeutics, by “inhibiting the inhibitor”; furthermore, any variations in tissue-specific (or tumor-specific) accumulation patterns of the antagonist vs CCL2 may lead to an unwanted increase in CCR2 stimulation where the inhibitor is not available. Overall, these issues suggest the need for even better CCR2-targeting compounds with higher affinity, longer residence times [51–61], broader tissue distribution, higher metabolic stability, and potentially entirely novel, scavenging-sparing pharmacological mechanisms.

Redundancy of the chemokine system is another factor that complicates the development of efficacious CCR2-targeting therapeutics and, in theory, provides a rationale for the development of multi-targeted inhibitors. In fact, some CCR2 inhibitors have comparable affinity for CCR5: for example, the former Merck clinical candidate **MK-0812** is equally potent on CCR2 and CCR5 [62,63] while the former ChemoCentryx clinical candidate **CCX-140** is CCR5-inert. The dual CCR2/CCR5 antagonism could be beneficial in some disease contexts [12,28,29,47,64,65] but detrimental in others. For example, in solid tumor immunology, both CCR2 and CCR5 contribute to the recruitment of MDSCs and macrophages, the key mediators of immunosuppression; however, CCR5 also drives the intratumoral recruitment of CD103+ dendritic cells which, through their effect on CD8+ effector T-cells, play a key role in both anti-tumor immunity and responsiveness to anti-PD-1 therapies [66,67]. Consequently, better pharmacological tools are needed to further interrogate the effect of CCR2 and CCR5 inhibition separately or collectively.

The *in vivo* pre-clinical analysis of CCR2 antagonists, and the translation of results from mouse models to human diseases, are often confounded by markedly lower antagonist affinity for mouse and other non-primate CCR2 compared to the human receptor [47]: this is the case, for example, for the ChemoCentryx candidate **CCX-140**. The known mouse-active CCR2 inhibitors almost universally hit mouse CCR5 as well, with only a few notable exceptions from the INCB compound series [68–70]. Thus, there is a critical need for CCR2-selective and dual compounds that are potent on both human and mouse CCR2 and have excellent pharmacokinetic properties, as tools for studying the pharmacological roles of CCR2 blockade in animal models of disease.

Many outstanding challenges in CCR2 inhibitor discovery and optimization can be addressed rationally, through structure-based drug design (SBDD). Indeed, atomic-level understanding of antagonist interaction with CCR2 and CCR5 may drive the optimization of compound pharmacokinetics (PK), potency, and the desired receptor and species selectivity. To aid such efforts and to complement the previously published structure of **maraviroc**-bound CCR5 [71], two structures of antagonist-bound CCR2 were solved in 2016 and 2019 [32,63]. More recently, the well-established physics-based SBDD approaches have been augmented by rapidly evolving artificial intelligence (AI) and deep-learning-based methods [72]. Nevertheless, receptor modeling, compound docking, and screening remain complex tasks involving numerous user-customizable parameters that may result in subjective outcomes. Accordingly, innovative frameworks must be developed and optimized to provide the best insight into rational SBDD for CCR2 and other GPCRs.

Here, using molecular docking and virtual compound library screening, we systematically identify the determinants of binding affinity and receptor and species selectivity for various chemotypes of orthosteric and allosteric antagonists of CCR2. The generated 3D models of receptor complexes with selective and dual-affinity antagonists, including the former clinical candidates **cenicriviroc** [41], and **CCX-140** [9] and the current clinical candidates **BMS-813160** [73,74] and **PF-4136309** [38], reveal hitherto unappreciated mechanisms of antagonist affinity and selectivity. Findings are further corroborated by insights from compound structure-activity relationship (SAR) series and rationally guided gain-of-function receptor mutagenesis.

Furthermore, to advance the SBDD efforts for CCR2, we reveal strong chemotype selectivity of the two available inactive-state experimental CCR2 structures and demonstrate the high predictive potential of a new deep-learning-based compound scoring function. Finally, we computationally generate new CCR2 models with high recognition capacity towards diverse chemotypes of orthosteric and allosteric antagonists, and establish the utility of limited-size, high-accuracy conformational ensembles for structure-based antagonist discovery.

Altogether, this work sheds light on novel strategies for creating highly selective CCR2 antagonists that leverage previously unexplored molecular interactions and offers new opportunities for rational compound design and optimization.

## Results

### CCR2 and CCR5 share structural principles when inhibited by orthosteric antagonists

To understand the conformational changes induced in the receptors by distinct ligands, we capitalized on the available X-ray and cryo-EM structures of CCR2 and CCR5 complexes with chemokine agonists or small molecule antagonists (**Fig. S1**). In CCR2, the orthosteric antagonist induces contraction of the binding pocket in the TM3-to-TM6/7 direction while simultaneously expanding the pocket in the perpendicular direction, thus increasing the TM1-to-TM7 distance to form an inter-helical, bilayer-facing gap (“fenestration”, **Fig. 1A-B**). The contraction of the TM3-to-TM6/7 distance and formation of the fenestration via TM1-TM7 separation similarly occurs in CCR5 (**Fig. 1C-E**). Both CCR2 antagonists that have been structurally characterized so far (**BMS-681** [32] and **MK-0812** [63]) protrude through this fenestration towards the membrane lipids. Interestingly, even though the gap is also present in CCR5 antagonist-bound structures [75,76], the antagonist maraviroc does not occupy it. This suggests that the TM1-to-TM7 expansion and the resulting formation of the fenestration are a common mechanism associated with inhibition of the two receptors by antagonists with diverse binding modes and chemotypes (**Fig. 1E**).

**Figure 1.**
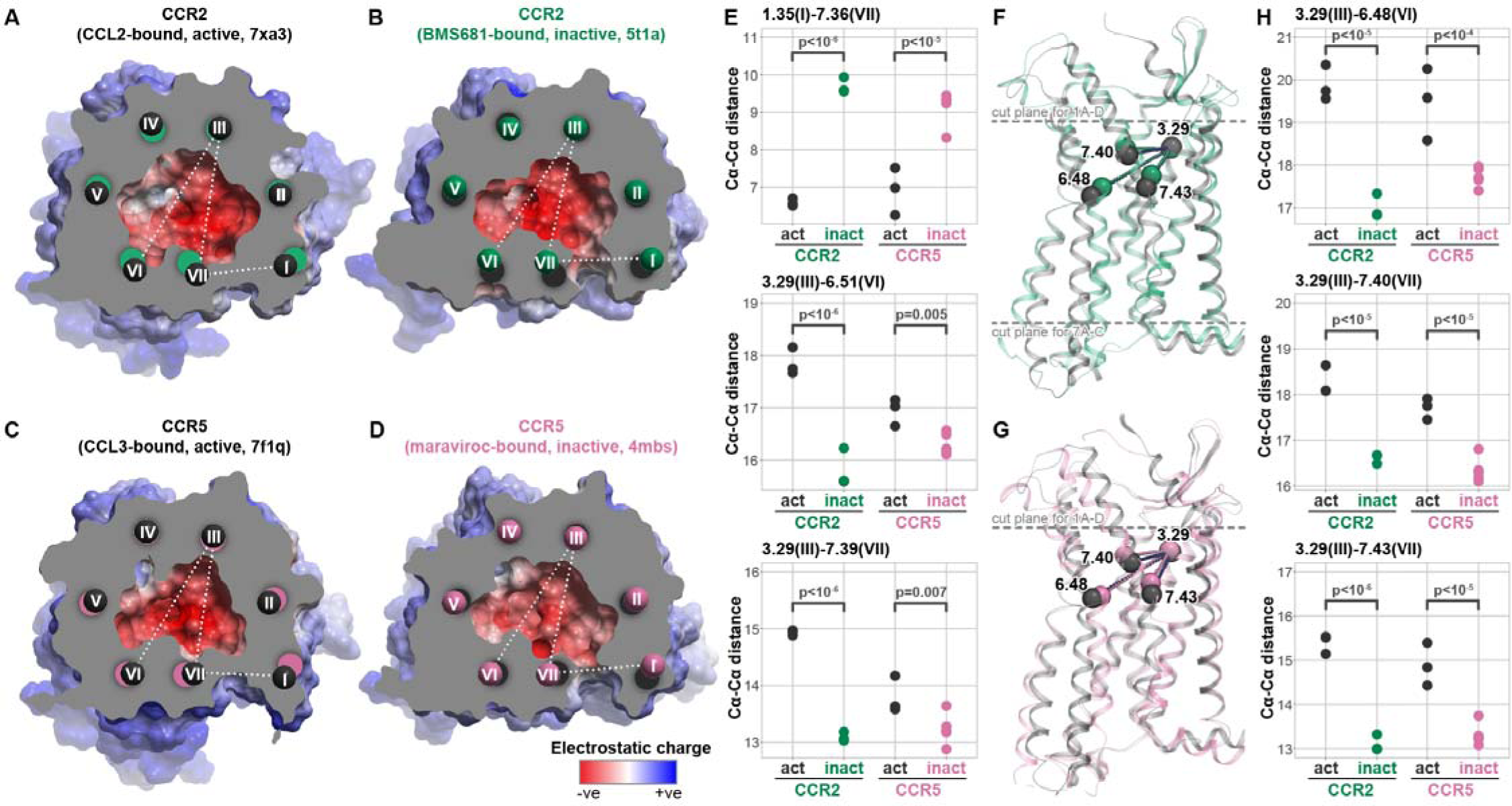
Shared structural principles of CCR2 and CCR5 inhibition by orthosteric antagonists. (**A-D**) Cross-sectional view of the orthosteric pockets of active (chemokine agonist-bound) (**A**) and inactive (antagonist-bound) (**B**) CCR2 in comparison to active (**C**) and antagonist-bound (**D**) CCR5, colored for electrostatic potential, demonstrate similar contraction of the TM6-to-TM3 and TM7-to-TM3 distances upon antagonist binding, with concurrent opening of the TM1-TM7 gap. The TM centroids are colored with a specific comparison: black - active, green - inactive CCR2, pink - inactive CCR5. (**E**) Cα-Cα distances for representative residue pairs from TM1, TM3, TM6, and TM7 measured across multiple active (agonist-bound) and inactive (antagonist-bound) structures of CCR2 and CCR5. (**F-G**) The subtle antagonism-associated rearrangements in the binding pocket translate into larger rearrangements of the transmembrane helices towards the middle of the bilayer and especially the intracellular side of both CCR2 (**F**) and CCR5 (**G**). Dashed lines indicate the placement of the cross-section planes for (**A-D**) as well as for Fig. 7A**-C**. (**H**) Cα-Cα distances between BW residue 3.29 (TM3) and representative residues in TMs 6 and 7 close to the middle of the bilayer, measured across multiple active (agonist-bound) and inactive (antagonist-bound) structures of CCR2 and CCR5.

Subtle rearrangements in the binding pocket translate into larger differences lower in the TM domain [77]. Specifically, in the agonist-bound state, both CCR2 and CCR5 demonstrate a large increase in the distance from the TM3 residues in the binding pocket (e.g. Ballesteros-Weinstein [78] (BW) position 3.29) to the middle of the TM helices 6 and 7 (e.g. BW 6.48, 7.40, 7.43) that indicate an outward motion of these helices and a vertical (i.e. perpendicular to the plane of the membrane) drop (**Fig. 1F-G**). Pairwise distances measured between these residues across available experimental structures of CCR2 and CCR5 indicate that the magnitude of change within the binding pocket is comparable between receptors (**Fig. 1E-H**). Collectively, these comparisons suggest a conserved mechanism for CCR2 and CCR5 orthosteric antagonism, whereby antagonists induce similar binding pocket conformations in the two receptors and prevent the subtle pocket expansion that leads to the downstream activation-associated helical rearrangements.

### Selected structures of CCR2 recognize known orthosteric antagonists in retrospective virtual screening

To assess the impact of these subtle binding pocket rearrangements on orthosteric antagonist binding to CCR2, we performed a retrospective virtual ligand screening (VLS) using a diverse library of 45 known CCR2 and CCR5 orthosteric antagonists (**Fig. S2-S4**) and a 54-fold larger set of property-matched decoys obtained from the DUD-E [79]. As with any VLS campaign, this involved (i) generation of multiple predicted poses for each compound in the binding pocket of the receptor, (ii) pose scoring and selection (where for each compound, one pose is selected from multiple generated), and (iii) compound scoring and ranking (where the comparison is performed across compounds, based on one selected pose for each). Screening was performed using the MolSoft ICM software suite [80] where (i) compound poses are efficiently generated using the receptor representation in the form of grid potential maps, after which (ii) pose selection and (iii) compound ranking are performed using one of three ICM scoring methodologies: full-atom (FA [81]), mean-force (MF [82]), or the recently implemented Radial-Topological Convolutional Neural Network (RTCNN [83,84]). An ideal score for stage (ii) robustly prioritizes the correct binding pose among multiple alternative predicted poses for that one compound, whereas an ideal score for stage (iii) provides a reliable comparison between different compounds in their correctly predicted binding poses. Success of VLS with each scoring scheme against each receptor pocket conformation was quantified as ROC AUC (area under the receiver operating characteristic curve [85]) which plots the number of true positives (known active compounds) against false positives in the top part of the compound list ordered by score. High ROC AUC (close to 100% on a 0%-100% scale) indicates that top-ranked compounds are indeed more likely to be potent binders towards the target. AUC of 50% indicates the lack of recognition, and AUC<50% means that the model is anti-predictive. To be practicable for prospective VLS of large databases, the model must place actives in the top 0.01%-1% of the list which corresponds to retrospective ROC AUCs as high as 99%, with initial (a.k.a. early, or top-of-the-list) recognition being especially important.

When using the conventional force-field-based FA method for both pose-selecting and ranking (referred to as the FA|FA scheme), we found that the orthosteric pockets of the antagonist-bound CCR2 structures (PDB 5t1a [32] and chains a, b of PDB 6gpx [63]), or their ensemble, provided reasonably strong early recognition of known CCR2 antagonists in VLS (**Fig. 2A**). By contrast, the pocket of the CCL2-bound active CCR2 structure (PDB 7xa3 [86]) had no predictive capacity at all (AUC=47.28%, **Fig. 2B**). This underlines the importance of the so-called “induced fit” and illustrates that even relatively subtle conformational changes in the binding pocket (such as those observed between the agonist- and antagonist-bound conformations of CCR2, see **Fig. 1**) may have a dramatic effect on the ability of the pocket to recognize compounds in VLS. The higher predictive capacity of antagonist-bound pockets in comparison to chemokine-bound pockets held for CCR5 conformers as well (**Fig. 2C-D**), although the differences were not as striking due to the small number of CCR5 orthosteric antagonists (10 for CCR5 vs 45 for CCR2) in the compound set.

**Figure 2.**
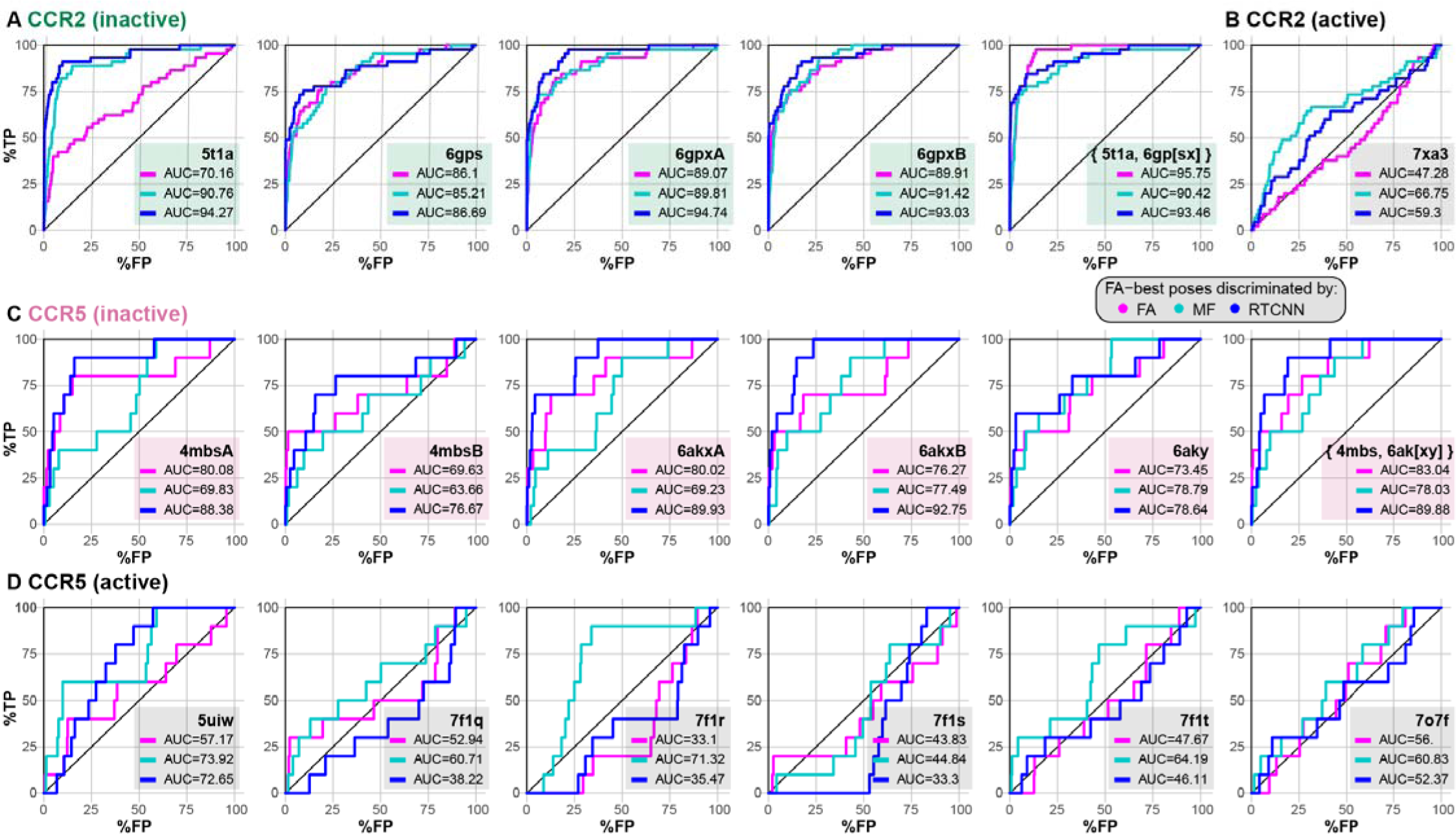
Selected structures of CCR2 recognize known orthosteric antagonists in retrospective virtual screening. (**A-B**) receiver operator characteristic (ROC) curves obtained by VLS of a diverse library of 45 known CCR2 orthosteric antagonists and a 54-fold larger set of property-matched DUDe decoys against the orthosteric pockets in inactive (antagonist-bound) (**A**) and active (chemokine agonist-bound) (**B**) CCR2 structure(s). Compound poses were selected using the full-atom score (FA); compounds were ranked by FA, mean-force (MF) or RTCNN score. (**C-D**) ROC curves obtained by VLS of a small diverse set 10 known CCR5 orthosteric antagonists and same inactives and decoys as in (**A-B**) against the orthosteric pockets in inactive (antagonist-bound) (**C**) and active (chemokine agonist-bound) (**D**) CCR5 structure(s).

Interestingly, we observed a remarkable improvement in antagonist recognition in the inactive CCR2 PDB 5t1a when compound ranking (iii) was performed by RTCNN instead of FA (**Fig. 2A**). Although not as striking, this compound ranking method also led to improvement in other CCR2 and even CCR5 structures (**Fig. 2A-B**). Ranking compounds with MF led to improved recognition in some but not all cases, and seldom outperformed RTCNN. The superior compound ranking performance of RTCNN over FA and MF suggested that this scoring function has a greater tolerance for induced fit and the associated small conformational rearrangements in the receptor pocket.

We also explored the impact of the three scoring functions when applied at the stage of compound pose selection (ii). When compound poses were selected by MF (i.e. MF|FA, MF|MF, MF|RTCNN scoring schemes), VLS performance deteriorated in almost all cases (**Fig. S5A-D**). However, selecting compound poses by RTCNN (RTCNN|FA, RTCNN|MF, RTCNN|RTCNN schemes) led to a similar or slightly improved VLS performance for both CCR2 and CCR5 (**Fig. S5E-H**), with notable differences being an improvement for the inactive CCR2 PDB 5t1a and worsened prediction, when ranked by FA, for active CCR5 structures (PDB 7f1[qrst], 7o7f). Based on these observations, the subsequent experiments were performed using only FA|FA or FA|RTCNN scoring schemes.

### CCR2 orthosteric pocket conformations are chemotype-selective

To understand the reason for consistently poorer FA|FA active/decoy discrimination by CCR2 PDB 5t1a compared to the other inactive CCR2 structures, we expanded our compound dataset by adding 357 published and patented Bristol-Myers-Squibb (BMS) compounds [87–91], and 113 patented Boehringer-Ingelheim (BI) compounds [92,93] of varying activity (**Supplemental Table 1**). Each set consisted of multiple representatives of a small number of structurally related chemotypes developed by the respective discovery program. Note that the original library already contained 21 published Merck CCR2 antagonists and their derivatives [26,94–99] (**Fig. S2, Supplemental Table 1**). We then performed chemotype-specific screens of the expanded library against all four experimental inactive state CCR2 structures.

Screening with this expanded compound set revealed that inactive orthosteric pockets best recognized compounds related to the ligands with which they were crystalized. For BMS compounds, PDB 5t1a (crystallized with **BMS-681**) provided very accurate discrimination but the performance of PDB 6gpx was much lower (**Fig. 3A**). Conversely, for Merck compounds and derivatives, PDB 6gpx (crystallized with **MK-0812**) provided almost ideal recognition whereas PDB 5t1a was virtually unable to discriminate them from decoys (**Fig. 3A**). Notably, both 5t1a and 6gpx had limited accuracy in BI compound recognition (**Fig. 3A**).

**Figure 3.**
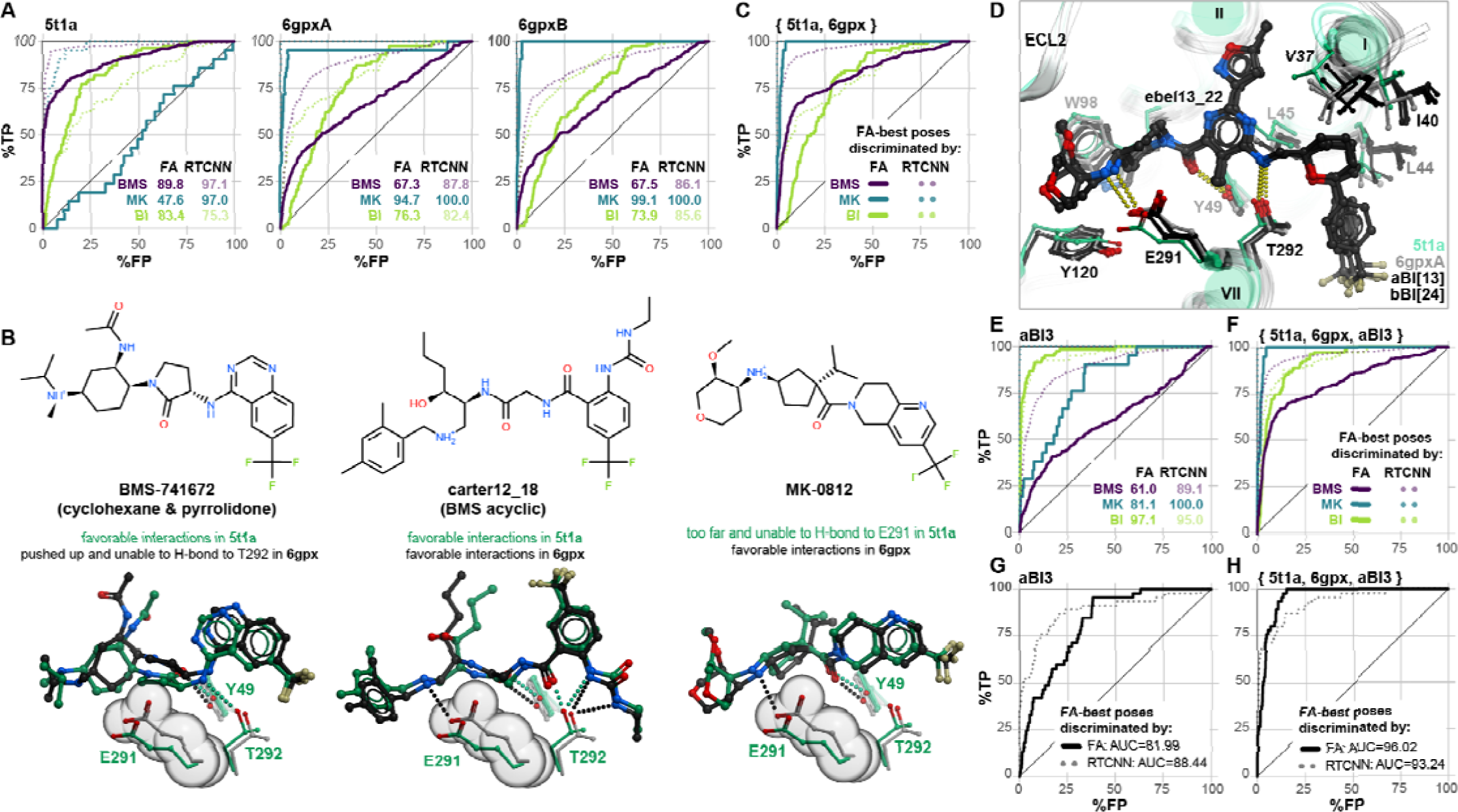
Chemotype selectivity of CCR2 orthosteric pocket conformations. (**A**) ROC curves obtained by VLS of expanded chemotype-specific libraries (330 BMS actives pIC_50_>7.5, 21 MK actives pIC_50_>7.5, 83 BI actives pIC_50_>7.5) and same inactives and decoys as in Fig. 2A**-B** against the orthosteric pockets in inactive (antagonist-bound) CCR2 structures PDB 5t1a and PDB 6gpx (chains A and B). Curves obtained using FA|FA and FA|RTCNN scoring schemes are shown in solid and dotted lines, respectively, with AUC values shown in the inset tables. (**B**) Differences in the conformation of E291^7.39^ between PDB 5t1a and PDB 6gpx explain chemotype-specific recognition. In PDB 6gpx, E291^7.39^ is oriented more towards the minor pocket (spheres) and creates steric interference with cyclic and pyrrolidone linkers of cyclic BMS compound while allowing for favorable docking of acyclic BMS compounds. By contrast, this orientation is required for Merck compounds that rely on it to form a salt-bridge with the charged amine. (**C**) ROC curves as in (**A**) but generated for an ensemble of experimental pocket conformations, PDB 5t1a and PDB 6gpx. (**D**) Representative CCR2 orthosteric pocket conformers generated through computational energy-based conformational optimization in the presence of **ebel13_22**. (**E-F**) ROC curves as in (**A**) using a representative computationally-generated BI-compatible conformer of 6gpx, aBI3 (**E**) or an ensemble of aBI3 and the experimental PDB 5t1a and PDB 6gpx (**F**). (**G-H**) ROC curves obtained by VLS of the initial 45 CCR2 orthosteric antagonist library and a 54-fold larger set of decoys against the computational generated conformer aBI3 (**G**) or the ensemble of aBI3, PDB 5t1a and PDB 6gpx (**H**) (compare to **Fig. 2A**).

We narrowed down the reasons for BMS vs Merck chemotype selectivity to a conformation of a single CCR2 residue, E291^7.39^: the elevated position of this residue in PDB 6gpx is favorable for the binding and salt bridge formation with the charged amines of the Merck compounds and acyclic BMS compounds, but prohibitive to the binding of cyclohexane- and pyrrolidone-containing BMS compounds, whereas the lowered position of E291^7.39^ in PDB 5t1a makes it compatible with most BMS compounds but disrupts the salt bridge critical for the Merck chemotypes (**Fig. 3B**). The ensemble of three experimental structures (BMS-selective PDB 5t1a and the two chains of Merck-selective PDB 6gpx[AB]) combined “the best of both worlds” and provided quality recognition of BMS and Merck compounds; however, it still largely failed to predict BI compounds (**Fig. 3C**).

To solve this problem, we applied computational energy-based conformational optimization to the pockets from PDB 6gpx in the presence of a docked **ebel13_22**, a potent BI CCR2 antagonist that encompassed most features of other compounds in the two BI series [92,93]. This step produced pocket conformers that were more compatible with BI compound binding, through widening of the TM1-TM7 gap and small changes in TM1, 2, and 7 residue conformations (**Fig. 3D**). The 6gpxA-derived conformer aBI3 demonstrated a much-improved recognition of BI compounds, at the expense of predictive capacity towards BMS and MK series (**Fig. 3E**), again emphasizing the impact of induced fit. Importantly, combining this conformer with experimental CCR2 PDBs 5t1a and 6gpx yielded an ensemble that was highly accurate towards all three chemotypes of CCR2 antagonists (**Fig. 3F**). The computationally generated conformer did not perform well in the original dataset of 45 actives vs decoys (**Fig. 3G**), because the set only contained one BI compound; however, again, its addition to the ensemble retained performance (**Fig. 3H**, compared to **Fig. 2A**). Thus through computational conformer generation, our screening methodology was expanded to recognize active compounds whose chemotype was unrepresented in available experimental structures.

The above observations related to the VLS strategy where both pose selection and compound ranking were performed using the physics-based FA score. Interestingly, chemotype sensitivity was much less of an issue when FA-best compound poses were discriminated by RTCNN: despite the induced fit and the observed conformational differences, all experimental structures and the computationally generated conformer aBI3 were predictive towards BMS and BI compound series (**Fig. 3A, C, E**). This illustrates that the AI-based RTCNN scoring function is more tolerant to small conformational inaccuracies in the binding pockets, and may provide superior discrimination of active compounds of multiple chemotypes in virtual screening.

### Structural determinants of CCR2 orthosteric antagonist affinity

Having confirmed the ability of CCR2 structures to recognize known antagonists in virtual screening, we next turned our attention to their predicted binding poses, to understand the molecular determinants for orthosteric antagonism. Notwithstanding their different chemotypes and scaffolds (**Fig. S4**), top-ranking poses of all CCR2 antagonists from the original set of 45 (**Fig. S2**) shared the overall binding geometry of **BMS-681** and **MK-0812** elucidated by crystallography [32,63]: they occupied the so-called minor subpocket of CCR2, packed against Trp98^2.60^, and protruded towards the bilayer through the inactive-state-specific TM1-TM7 fenestration (**Fig. 1B**). **Fig. 4A-**I illustrates these principles for four diverse clinically relevant orthosteric antagonists: the CCR2-selective antagonist **PF-4136309** [68] (**Fig. 4A, E**) and the dual-selectivity CCR2/CCR5 antagonists **BMS-813160** [73] (**Fig. 4B, F**), **cenicriviroc** (Takeda) [100] (**Fig. 4C, G**), and **qin18_7** (Boehringer Ingelheim) [101] (**Fig. 4D, I**). Despite scaffold variations, all examined compounds employed an amide linker or isostere to hydrogen-bond with Y49^1.39^ and/or T292^7.40^ in the TM1-TM7 tunnel (**Fig. 4J-M**) (residues W98^2.60^, Y49^1.39^, and T292^7.40^ are all conserved between CCR2 and CCR5). In addition to these universal interactions, many compounds formed a salt-bridge or hydrogen-bond with CCR2 E291^7.39^ (**Fig. 4M**) and reached the cavity under P192^45.52^ of ECL2 (**Fig. 4E-M**). We also identified some semi-unique binding modes and interactions: for example, **PF-4136309** extended towards the major subpocket and hydrogen-bonded to T179^4.64^ (**Fig. 4J**), whereas **cenicriviroc** utilized its large flexible butoxy-ethoxy-phenyl moiety to bind extrahelically at the interface with membrane lipids, in the TM1-TM7 groove (**Fig. 4H**). The **PF-4136309** mode of binding (utilizing the major subpocket) was shared by **INCB-3284** and **INCB-3344** [69,102] (**Supplemental Data 1**). Surprisingly, we also found that the extrahelical binding mode of **cenicriviroc** was shared by several other antagonists created by Takeda (**TAK-779** and derivatives [103,104], contrasting earlier predictions [105–109]) and BI [93] (**Supplemental Data 1**). These conserved and unique binding determinants may provide insight for future structure-guided optimization of CCR2 antagonist properties.

**Figure 4.**
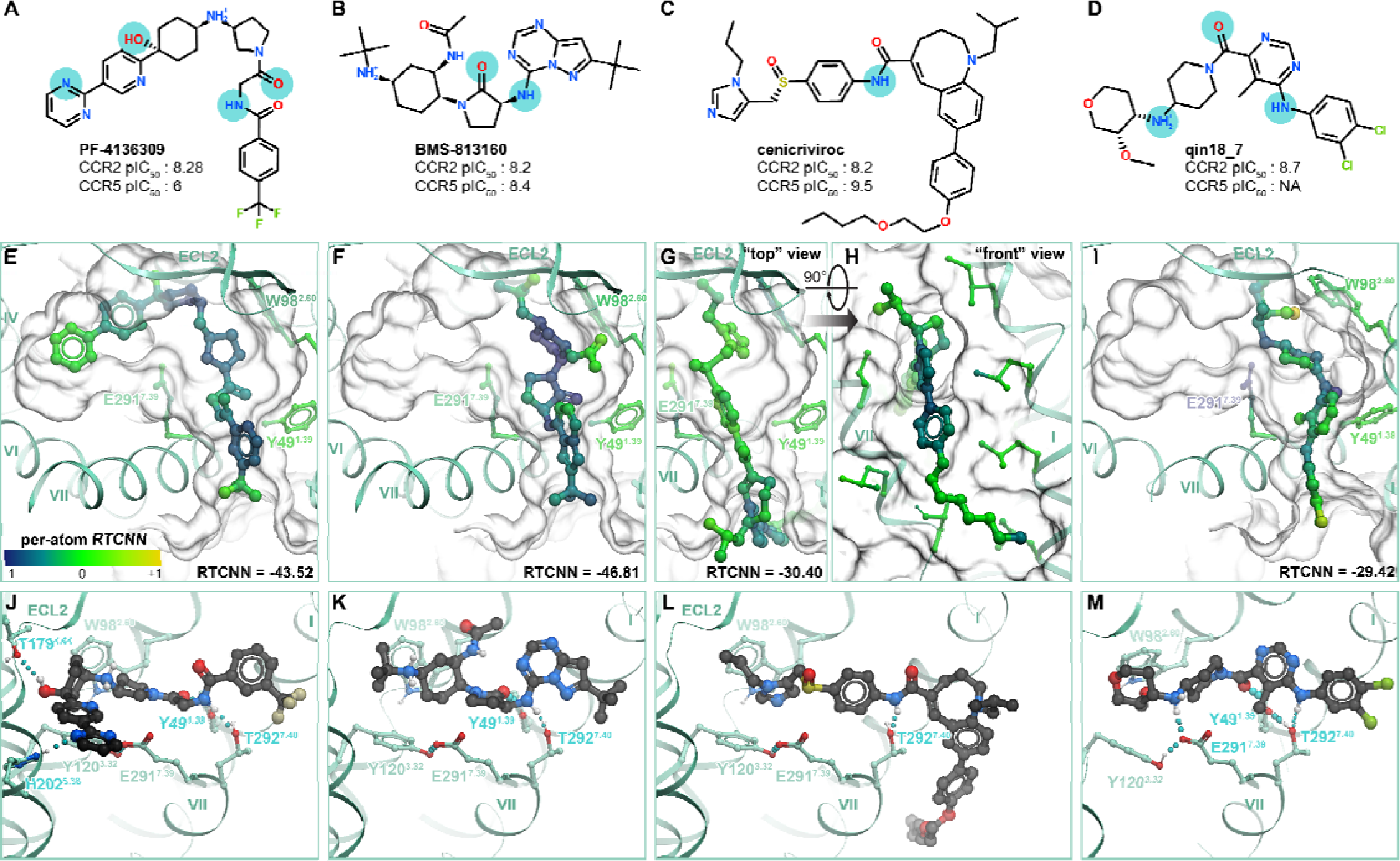
Structural determinants of antagonist binding to the orthosteric pocket of CCR2. (**A-D**) Chemical structures of clinically relevant orthosteric antagonists **PF-4136309** (**A**), **BMS-813160** (**B**), **cenicriviroc** (**C**), and **qin18_7** (**D**) and their experimentally measured potencies (expressed as pIC_50_) for human CCR2 and CCR5. Cyan circles highlight atoms predicted to be involved in key polar interactions. (**E-I**) Predicted binding poses of compounds in (**A-D**) to human CCR2, viewed perpendicularly to the plane of the membrane from the extracellular side (**E**, **F**, **G**, **I**) or along the plane of the membrane (**H**). Compound atoms are colored for their contribution to the RTCNN score. Key interacting receptor residues are depicted as sticks, the surface of the binding pocket is shown as a transparent mesh. (**J-M**) Predicted polar interaction networks for the compounds from (**A**-**D**) in the binding pocket of CCR2. Hydrogen bonds are shown as cyan dotted lines, receptor amino-acid residues involved in hydrogen bonding interaction are labeled in cyan.

The predicted antagonist binding poses and associated docking scores provided a way to rationalize “activity cliffs” and divergent sensitivities of previously tested compounds to receptor mutations. “Activity cliffs” are pairs of related compounds where minute changes to a compound’s chemical structure drastically alter target affinity [110,111]. For instance, the pair of BMS compounds **cherney10_21** and **cherney10_20** are stereoisomers, with a chiral carbon in the pyrrolidone’s 3’ position accounting for a ∼100x loss in activity towards CCR2 [112] (**Fig. 5A**). Predicted compound poses indicate that loss of hydrogen bonding to Y49^1.39^ may account for this reduction (**Fig. 5B-C**): this emphasizes the role of this hydrogen bond as one of the key affinity determinants. In another instance, the BI compound **ebel11_9** differs from **ebel11_17** by a change from 3,4-dichlorobenzyl to a 2-chlorobenzyl substituent, accounting for a ∼1000x reduction in activity [92] (**Fig. 5D**). Docking predicted that the 2’ chlorine introduces steric clashes with residues in TM1 and TM7 (**Fig. 5E-F**), coincident with a loss in RTCNN score: this illustrates the importance of precise steric fit within the TM1-TM7 fenestration for compound affinity.

**Figure 5.**
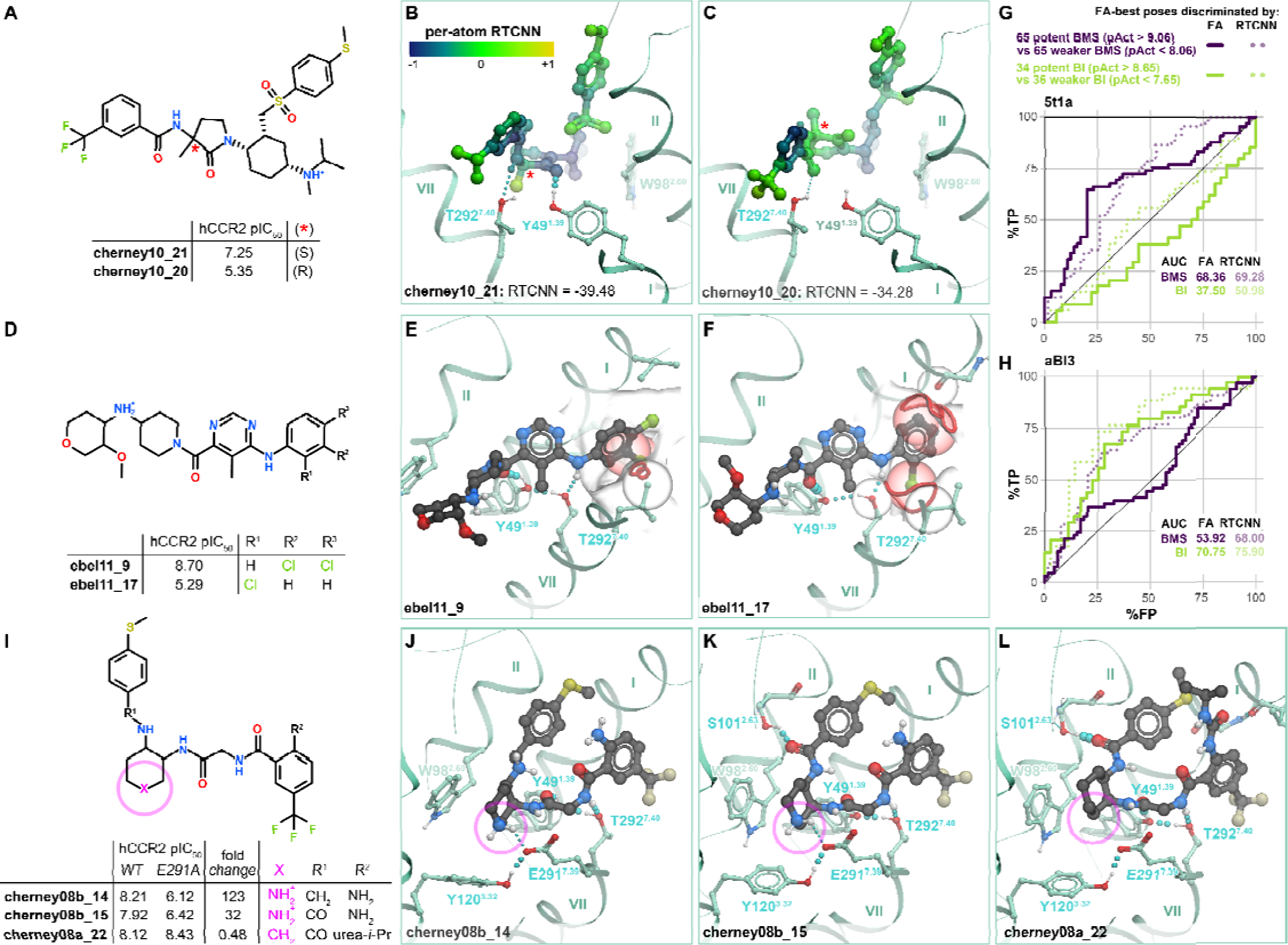
Models explain affinity differences within orthosteric compound series. (**A**) Chemical structures and CCR2 inhibitory potencies of stereoisomers **cherney10_21** and **cherney10_20**. Red asterisk indicates the stereoisomers’ chiral center. (**B-C**) Predicted binding poses of **cherney10_21** (**B**) and **cherney10_20** (**C**) demonstrate the steric differences and the loss of hydrogen bonding with CCR2 Y49^1.39^ in **cherney10_20**. Compound atoms are colored by their contribution to the RTCNN score with a notable difference at the pyrrolidone oxygen, consistent hydrogen bond loss and the aggregate RTCNN score. (**D**) Chemical structures and CCR2 inhibitory potencies for compounds **ebel11_9** and **ebel11_17**. with the (**E-F**) Predicted binding poses of **ebel11_9** (**E**) and **ebel11_17** (**F**) in the binding pocket of CCR2 demonstrate how rearrangement of phenyl substituents introduces steric collisions (clashes) with receptor pocket residues. Clashes are depicted as red rings demarking the spatial intersections between the receptor atoms (white mesh) and compound atoms (red spheres). Hydrogen bonding residues are labeled in cyan. (**G-H**) ROC curves obtained by challenging PDB 5t1a (**G**) or the computationally generated BI-compatible conformer aBI3 (**H**) with distinguishing potent compounds of either BMS (65 compounds, pIC_50_>9.06) or BI (34 compounds, pIC_50_>8.65) chemotypes from an approximately equal number of less-potent compounds of the corresponding chemotype (BMS: 65 compounds, pIC_50_<8.06, BMS: 36 compounds, pIC_50_<7.65). (**I**) 2D chemical structures and inhibitory potencies towards WT CCR2 and CCR2(E291^7.39^A) for the series of CCR2 antagonists **cherney08b_14**, **cherney08b_15**, and **cherney08a_22**. (**J-L**) Predicted poses of **cherney08b_14** (**J**), **cherney08b_15** (**K**), and **cherney08a_22** (**L**) in the binding pocket CCR2, viewed from the extracellular side. Hydrogen bonds are shown as cyan dotted lines. Residues that participate in hydrogen-bonding with the compound are labeled in cyan. The variable 3’ position of the cyclohexane is circled in pink. The top portion TM7 and all of TM6 are hidden for clarity.

Despite successes with individual activity cliffs, we noticed that neither the FA|FA nor FA|RTCNN scoring scheme could accurately distinguish potent actives from weaker actives or inactive compounds *of the same chemotype* when applied in the virtual screening mode. Specifically, when the BMS-compatible CCR2 pocket conformer PDB 5t1a was challenged with discriminating 65 potent BMS antagonists (pActivity>9.06) from an equal number of weaker BMS actives and inactives (pActivity<8.06), the ROC AUC was only 68.36% and 69.28% for the FA|FA and FA|RTCNN scoring schemes, respectively, with even the early recognition being unsatisfactory (**Fig. 5G**). Similarly, when the computationally-generated BI-compatible CCR2 pocket conformer aBI3 was presented with 34 BI potent actives (pActivity>8.65) and an approximately equal number of weaker BI actives and inactives (pActivity<7.65), the respective ROC AUCs were only 70.75% and 75.9% (**Fig. 5H**). Consistent with chemotype selectivity of these two conformers (**Fig. 3**), the screening performance was even lower when tested across (PDB 5t1a tested with BI compounds and BI-compatible conformer aBI3 tested with BMS compounds, **Fig. 5G-H**). Because neither FA nor RTCNN scoring functions were designed or trained to predict binding affinities, their observed inability to accurately distinguish potent actives from weaker actives of the same chemotype is unsurprising. Nevertheless, the chemotype-selective conformers provided some positive recognition towards their cognate compound chemotypes even in this challenging scenario.

Finally, the models provided a way to rationalize variations in compound sensitivity to mutations in the receptor binding pocket. For example, BMS reported structurally related compounds **cherney08b_14**, **cherney08b_15**, and **cherney08a_22**, which differ by the presence of a secondary amine inside the cyclical scaffold (**cherney08a_22** has a cyclohexane ring whereas **cherney08b_14,15** both have a piperidine ring, **Fig. 5I**). Our models suggest that the secondary amine in **cherney08b_14** makes an important salt bridge with CCR2 E291^7.39^, and one of only three hydrogen bonds (the other two being to Y49^1.39^ and T292^7.40^) that the compound makes in the binding pocket (**Fig. 5J**); this explains the >100-fold loss [113] in **cherney08b_14** potency towards the CCR2(E291^7.39^A) mutant compared to WT CCR2. For the piperidine-bearing compound **cherney08b_15**, which forms an additional (fourth) hydrogen bond in the pocket, to CCR2 S101^2.63^ (**Fig. 5K**), the E291^7.39^A mutation causes only a 32-fold potency loss. Notably, the replacement of piperidine by a cyclohexane, as in **cherney08a_22**, makes the compound completely insensitive [114] to the E291^7.39^A mutation: this is because it no longer relies on the hydrogen bond or a salt bridge to E291^7.39^ for affinity (**Fig. 5L**). By contrast, the T292^7.40^A mutation causes ∼40-fold loss in potency for **cherney08a_22** [114] and is detrimental for many other compounds [109,115], consistent with the predicted role of this residue as a universal hydrogen-bonding anchor for CCR2 orthosteric antagonism.

Taken together, these findings illustrate that the predicted docking poses and complex geometries can guide structure-based compound optimization. Additionally, the RTCNN scoring scheme enables a straightforward calculation of individual atom contributions to the overall score. Mapping these per-atom contributions onto the predicted compound poses helps identify the most significant/favorable molecular interactions as well as the interactions in need of optimization. For example, favorable RTCNN scores were associated with the conserved binding affinity determinants: packing with W98^2.60^, tucking under ECL2, and accessing the TM1-TM7 fenestration (**Fig. 4E-I**). Similarly, depicting per-atom RTCNN scores on the **cherney10_21/20** stereoisomers captured how the lost hydrogen bond negatively impacts the score (**Fig. 5A**).

### Structural determinants of CCR2/CCR5 orthosteric antagonist selectivity

Many but not all CCR2 antagonists share a comparable affinity to CCR5 [4,28,73,116]. The two receptors have overlapping but distinct functions in chemokine biology [66], making this dual selectivity desirable in some contexts [64,65] but not others [18,37,38,117]. Therefore, it is important to understand what features distinguish CCR2-selective antagonists from dual-affinity ones. Additionally, CCR2 is a GPCR with profound differences between species: a structural understanding of features defining species selectivity is necessary to guide and rationalize the results of animal (e.g. mouse) experiments with CCR2 antagonists. We thus sought to understand the determinants of orthosteric antagonist selectivity between human CCR2 (hCCR2), human CCR5 (hCCR5) and mouse CCR2 (mCCR2).

A hCCR2-vs-hCCR5 comparison of amino-acid residues lining the binding pocket revealed that most of the substitutions are located in the major subpocket and involve residues in TM5: N199^5.35^K, H202^5.38^Q, R206^5.42^I. N199^5.35^ and H202^5.38^ are also found as K and Q in mCCR2, whereas R206^5.42^ is conserved between human and mouse (**Fig. 6A-B**). The minor subpocket harbors only a single substitution with hCCR2 S101^2.63^ replaced by Y89^2.63^ in hCCR5; this residue is also tyrosine in mCCR2 (**Fig. 6A**). Finally, bridging the two subpockets, hCCR2 H121^3.33^ and P192^45.52(ECL2)^ are replaced by F109^3.33^ and S180^45.52(ECL2)^, respectively, in hCCR5, but are conserved in mCCR2 (**Fig. 6A**).

**Figure 6.**
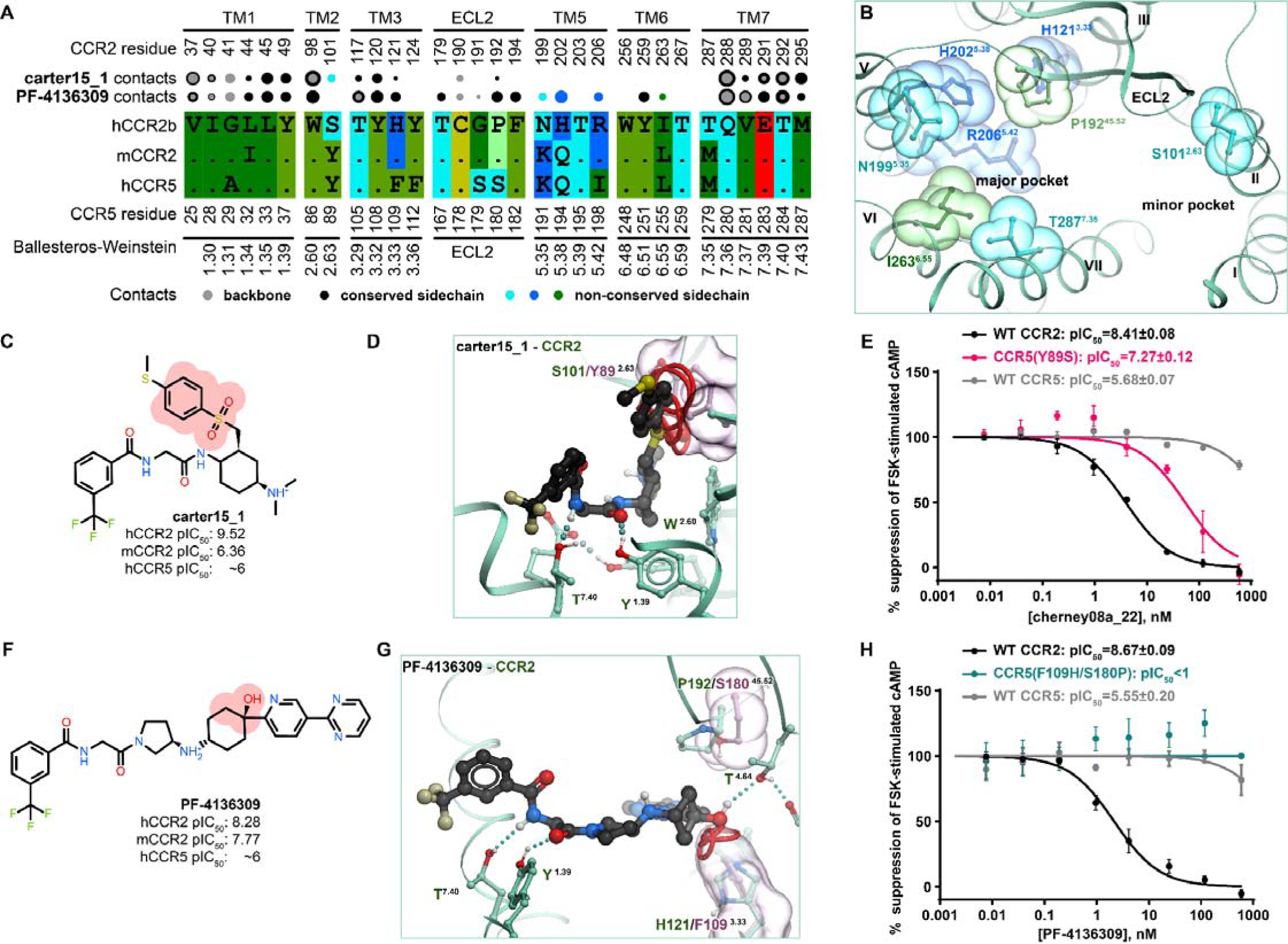
Structural determinants of CCR2/CCR5 orthosteric antagonist selectivity. (**A**) Sequence alignment and the map of the predicted contacts between the compounds and receptor amino-acid-residues for the hCCR2-selective minor-pocket binder **carter15_1** and the [hm]CCR2-selective and-minor pocket binder **PF-4136309.** Interacting non-conserved residues are shown in color. major- (**B**) The binding pocket of hCCR2, viewed across the plane of the membrane from the extracellular side, with residues that are not conserved in hCCR5 and/or mCCR2 shown as spheres and colored as in (**A**). (**C**) Chemical structure of hCCR2-selective antagonist **carter15_1** and its experimentally measured p tencies against hCCR2, hCCR5, and mCCR2. Highlighted with red circles are the atoms of the methylthio-phenyl-sulfonyl moiety predicted to collide with Y89^2.63^ of hCCR5 and Y114^2.63^ of mCCR2, but not with S101^2.63^ of hCCR2. (**D**) Superimposition of the docked pose of **carter15_1** in hCCR2 onto hCCR5. Emerging steric hindrances are highlighted between the methylthio-phenyl-sulfonyl moiety (red spheres) and Y89^2.63^ (pink surface). (**E**) Inhibition of chemokine-induced suppression of forskolin-stimulated cAMP by increasing concentrations of **cherney08a_22** in HeLa cells transiently expressing WT CCR5, WT CCR2, or CCR5(Y89^2.63^S). cAMP was measured via the BRET-based CAMYEL (cAMP sensor using YFP-Epac-RLuc) biosensor assay for detection of Gai-induced suppression of cAMP. Data was normalized to 0%-100% where 0% and 100% correspond to non-chemokine-stimulated and chemokine-stimulated, antagonist-free samples, respectively, in the same experiment. Error bars represent ±S.E.M. from n=3 independent experiments performed on different days, each with 2 technical replicates. Data was collated and fitted to a one-parameter non-linear regression model (log[inhibitor] vs. normalized response) in GraphPad Prism. (**F**) Chemical structure of [hm]CCR2-selective antagonist **PF-4136309** and its experimentally measured potency against hCCR2, hCCR5, and mCCR2. Highlighted with red circles is the methyl-cyclohexyl moiety predicted to collide with F109^3.33^ but not H121^3.33^. (**G**) Predicted hydrogen-bonding between **PF-4136309** and hCCR5 T167^4.64^ (corresponds to hCCR2 T179^4.46^) may be blocked by hCCR5 S180^45.54(ECL2)^ and F109^3.33^ (correspond to hCCR2 P192^45.52(ECL2)^ and H121^3.33^, respectively) to abrogate compound binding to hCCR5. (**H**) Same assay as in (**E**) evaluating interaction of CCR5(F109H/S180P) with **PF-4136309.**

Consistent with these observations, dual-selective (hCCR2/hCCR5) antagonists are predicted to bind exclusively within the minor subpocket (**Fig. 4B-D, F-I**) and have no, small, and/or flexible substituents in proximity of hCCR2 S101^2.63^ (e.g. no substituent in **cenicriviroc, Fig. 4C,G**; small substituents in **BMS-681** and **BMS-813160**, **Fig. 4B,F**). In fact, a bulky substituent next to S101^2.63^ appears sufficient for hCCR2 selectivity: it does not compromise compound affinity to hCCR2 but creates an unfavorable steric interference with Y89^2.63^ in hCCR5 and Y114^2.63^ in mCCR2. This is exemplified by **carter15_1** (**Fig. 6C**) where the steric clash between the compound methylthio-phenyl-sulfonyl moiety and hCCR5 Y89^2.63^ (**Fig. 6D**) (or Y114^2.63^ in mCCR2) abrogates binding to hCCR5 and mCCR2. Another example is **cherney08a_22** (also known as **BMS CCR2 22**, **Fig. 5I**). In accordance with this selectivity mechanism, we hypothesized that mutating hCCR5 Y89^2.63^ to match hCCR2 S101^2.63^ would permit hCCR5 binding to previously hCCR2-selective **carter15_1** and **cherney08a_22**. The hCCR5(Y89^2.63^S) mutant that we generated remained responsive to CCL5 (**Fig. S6**) and could indeed be inhibited by **cherney08a_22** (**Fig. 6E**), in contrast to WT hCCR5. This gain-of-function experiment supports the role of BW position 2.63 as a determinant of selectivity towards hCCR2 for this class of antagonists.

Notably, although a bulky substituent next to S101^2.63^ is *sufficient* for hCCR2 selectivity, it is not *necessary*. Compounds with medium-sized substituents such as acetamide can still demonstrate a wide range of activities towards CCR5, exemplified by **BMS-813160** (**Fig. 4B, F**) and the series of related compounds in Yang et al. [118]. Models suggest that the observed variations in hCCR5 affinity may be attributed to Y89^2.63^-mediated alteration of compound interactions with the cavity under the receptor’s ECL2: while dual hCCR2/hCCR5 antagonists employ favorable hydrophobic interactions in the TM3-ECL2 subpocket with their invariable tert-butyl substituent (e.g. **BMS-813160** or **yang21_6, Fig. S7A-E**), hCCR2-selective compounds exclusively rely on hydrogen bonding with the only proximal donor, the backbone carbonyl of C190^45.50^ (e.g. **yang21_5a**, **Fig. S7F-I**), located in hCCR2 ECL2 and sterically blocked by hCCR5 Y89^2.63^.

Finally, CCR2 selectivity can also be achieved in compounds that lack interactions with S101^2.63^ entirely and instead extend into the major subpocket of the receptor: this is exemplified by **PF-4136309** (**Fig. 4A,E**), **INCB-3284**, and **INCB-3344** (**Supplemental Data 1**). Extending into the major subpocket allows these compounds to interact with H121^3.33^, P192^45.52(ECL2)^, and sometimes R206^5.42^ (**Fig. 4E, J, Fig. 6G**) which are all conserved in mCCR2 but not in hCCR5 (**Fig. 6A**), and thus achieve selectivity against hCCR5 while remaining mouse-active. For **PF-4136309,** H121^3.33^ and P192^45.52(ECL2)^ gate compound access and hydrogen bonding to the conserved T179^4.64^ (**Fig. 4E**); superimposing the predicted hCCR2-bound pose of **PF-4136309** with hCCR5 suggests that S180^45.52^ and F109^3.33^ create substantial hindrances that could impede compound access to T167^4.64^ of hCCR5 thus prohibiting the interaction (**Fig. 6G**). As this mechanism was also proposed for other CCR2-selective compounds [63], we chose to probe its role in hCCR2 selectivity of **PF-4136309**. The double mutant of hCCR5, F109^3.33^H/S180^45.52(ECL2)^P, was slightly more responsive to CCL5 than WT hCCR5 (**Fig. S6)** but did not gain sensitivity to **PF-4136309 (Fig. 6H**), thus disproving the role of H121^3.33^ and P192^45.52(ECL2)^ in [hm]CCR2 selectivity vs CCR5 selectivity vs CCR5 for at least this one compound (it may still be the mechanism for e.g. **INCB-3284** and **INCB-3344**). Through process of elimination, we conclude that for **PF-4136309**, the selectivity determinant is likely R206^5.42^ (R219^5.42^ in mCCR2, I198^5.42^ in hCCR5). Because I198^5.42^ is important for chemokine binding to hCCR5, we decided against generating the hCCR5 I198^5.42^R mutant, as its effects on competitive blockade of chemokine binding to hCCR5 would be difficult to deconvolute.

### Only inactive holo structure of CCR2 recognizes allosteric antagonists in virtual screening

The intracellular side of CCR2 harbors a druggable pocket which is targeted by numerous known allosteric CCR2 antagonists [30,32]. Of the four available inactive-state CCR2 structures, only one (PDB 5t1a [32], **Fig. 7A**) features an allosteric antagonist, CCR2-RA-[*R*] [119], occupying this pocket. The remaining inactive CCR2 structures (PDB 6gp[sx] [63]) also have a cavity in the corresponding location; however, it is substantially altered by thermo-stabilizing mutations introduced to aid structure determination [63], and differs from the holo-cavity in PDB 5t1a even after the mutations are reverted to WT (**Fig. 7B**). Moreover, the unresolved C-terminal helix 8 in PDB 6gpxB creates additional void space (**Fig. 7B**). Finally, in the chemokine- and G protein-bound CCR2 PDB 7xa3, the activation-associated helical rearrangements (**Fig. 1F**) lead to a complete collapse of the cavity and the formation of the G protein interface instead (**Fig. 7C**).

**Figure 7.**
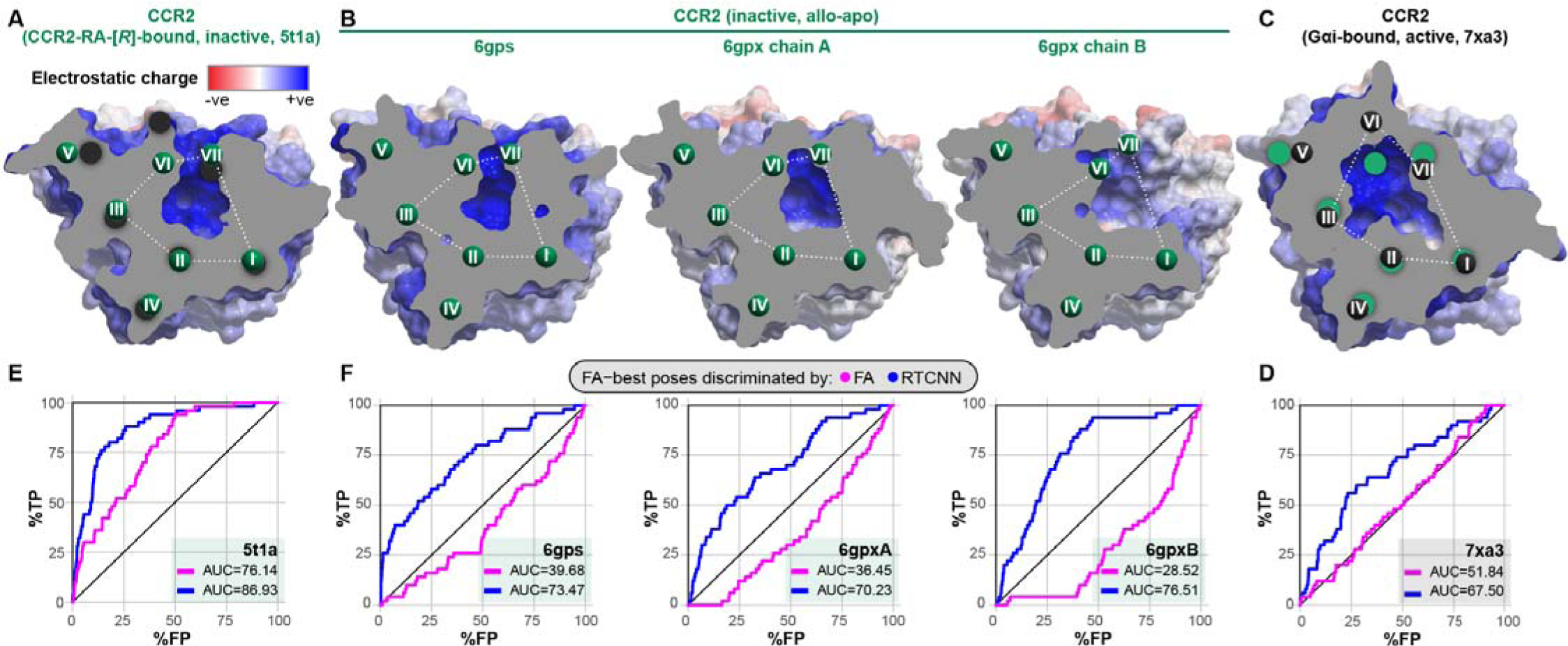
Appearance and retrospective VLS performance of the intracellular allosteric pockets in the available experimental structures of CCR2. (**A-C**) Cross-sectional view of the allosteric pockets of inactive antagonist-bound (**A**), inactive allo-apo **B**), and active G protein-bound (**C**) CCR2, colored for electrostatic potential and viewed perpendicularly to the plane of the membrane from the intracellular side. The position of the cross-section plane is shown in Fig. 1F. The TM centroids are colored black for the active conformation and green for the inactive CCR2 PDB 5t1a. (**D-F**) ROC curves obtained by VLS of a diverse library of 50 known CCR2 allosteric antagonists and a 59-fold larger set of property-matched DUDe decoy compounds (2,950 compounds) against the respective pockets in (**A-C**). Compound poses were selected using the full-atom score (FA); compounds were ranked by either FA or RTCNN score.

To understand the consequences of these conformational differences to antagonist recognition, we challenged these allosteric pocket conformations with a task of discriminating 50 known diverse allosteric antagonists (**Fig. S3-S4**) from 59x as many property-matched decoys from the DUD-E in retrospective VLS. As expected, the G protein-bound CCR2 conformation (PDB 7xa3) completely failed to recognize active compounds among decoys using the FA|FA scoring scheme (**Fig. 7D**). Importantly, among the inactive-state structures, only the holo structure (PD 5t1a) demonstrated appreciable ROC AUC (76.14%, **Fig. 7E**), with the apo structures being anti-predictive (**Fig. 7F**): this suggests that pocket shape and induced fit are even more critical for allosteric than for orthosteric antagonists. As with orthosteric VLS, we observed that compound ranking by RTCNN (the FA|RTCNN scheme) leads to an apparent improvement in screening accuracy. However, this improvement was not associated with correct docking poses or interactions: such interactions were unachievable in the active-state CCR2 PDB 7xa3 or in the inactive CCR2 PDB 6gpxB that lacks the entire helix 8 and all polar hydrogens important for antagonist recognition. Taken together, this suggests that RTCNN scoring may become a source of false positives in screening through *excessive* tolerance to conformational variability, and that FA|FA scoring should not be dismissed. We therefore again sought ways to optimize the FA|FA recognition by PDB 5t1a towards multiple chemotypes of allosteric CCR2 antagonists.

### CCR2 allosteric pocket conformations are antagonist-chemotype-selective

Guided by the insights from the CCR2 orthosteric antagonist VLS (**Fig. 3**), we probed the ability of the only predictive CCR2 allosteric pocket conformer (PDB 5t1a, **Fig. 7A**) to selectively recognize compounds of individual chemotypes. For this purpose, we assembled a larger library containing 113 AstraZeneca and related allosteric CCR2 antagonists (AZ [120–123]), 52 UCB Pharma antagonists [119,124], 93 Johnson & Johnson antagonists (JNJ, [125–127]), and 79 GlaxoSmithKline antagonists (GSK [128]) from the literature, as well as 50 published [129] and patented [130] BMS compounds, and 40 patented ChemoCentryx (CCX) compounds [131–133]. The allosteric pocket from CCR2 PDB 5t1a was then challenged with recognizing active compounds of individual chemotypes against inactives (from all chemotypes) and the original set of 2950 decoys.

As was the case with the orthosteric holo pockets, substantial biases were observed. The allosteric pocket of PDB 5t1a provided excellent recognition of AZ, UCB, and BMS compounds; however, the recognition of JNJ, CCX, and GSK compounds was very poor (**Fig. 8A**). The superior performance of PDB 5t1a towards the UCB chemotype is not surprising because the structure was solved with a representative of that chemotype, the antagonist **CCR2-RA-[*R*]** (**Fig. S3**). The shape of the pocket also appears suitable for BMS and AZ compounds. However, GSK compounds and **CCX-140** [133–135] appeared to be too large, thus scoring poorer than many decoy compounds or inactives from other, better fitting chemotypes. Nevertheless, the predicted binding poses for at least some compounds of these poorly performing chemotypes were conformationally feasible and consistent with their published SAR. For instance, for **JNJ-27141491**, the 5-ester group essential for potency [126] was predicted to engage in hydrogen-bonding, while for **CCX-140**, hydrogen bonds were formed with its sulfonamide.

**Figure 8.**
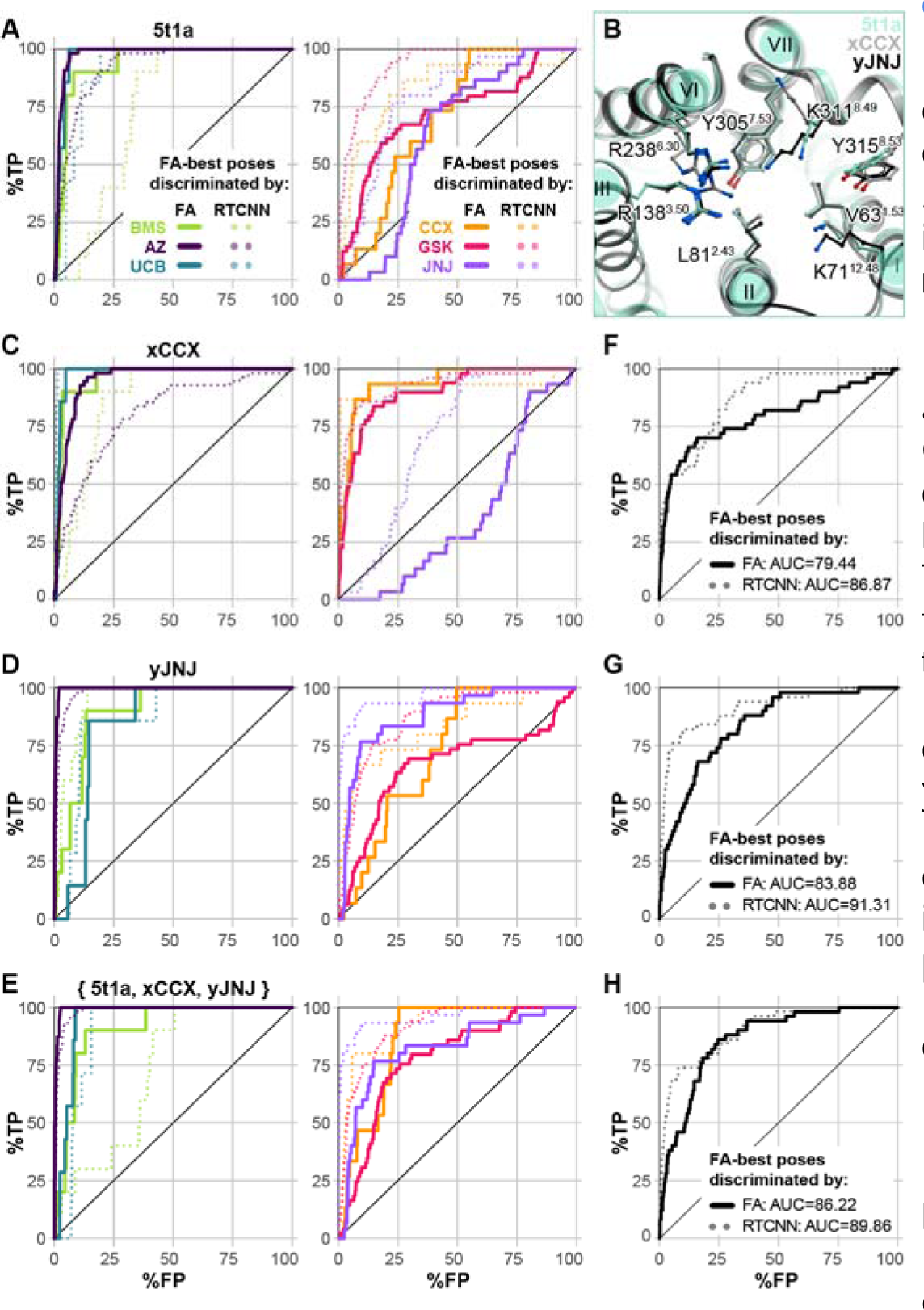
Allosteric recognition by 5t1a is also chemotype-specific. (**A**) ROC curves obtained by VLS of expanded chemotype-specific libraries and a pooled set consisting of all-chemotype inactives and the original 2950 decoys (Fig. 7D**-F**), against PDB 5t1a. For the individual color-coded curves, the positive parts of the respective libraries consisted of 10 BMS actives (pIC_50_>7.0), 55 AZ actives (pIC_50_>7.0), 7 UCB actives (pIC_50_>6.5), 15 CCX actives (pIC_50_>7.0), 49 GSK actives (pIC_50_>7.0), or 30 JNJ actives (pIC_50_>7.0). Curve style indicates whether FA-best poses were discriminated by FA (FA|FA scoring scheme, solid lines) or RTCNN (FA|RTCNN scoring scheme, dots). The two charts partition chemotypes recognized favorably by PDB 5t1a (BMS, AZ, and UCB) from those recognized poorly (CCX, GSK, and JNJ). (**B**) Superimposition of PDB 5t1a and two computationally generated conformers, xCCX and yJNJ, minimized in the context of restrained **CCX-140** (xCCX) or **JNJ-27141491** (yJNJ) and chosen for their capacity to recognize their corresponding chemotypes in VLS. Structures are viewed perpendicular to the plane of the membrane from the intracellular side. (**C-E**) ROC curves obtained by VLS of the same compound libraries as in (**A**) against xCCX (**C**), yJNJ (**D**), or the ensemble of xCCX, yJNJ, and PDB 5t1a (**E**). (**F-H**) ROC curves obtained by VLS of the original library of 50 diverse allosteric antagonists and 2,950 DUDe decoys against xCCX (**F),** yJNJ (**G**), or an ensemble comprising xCCX, yJNJ, and PDB 5t1a (**H**).

Because the pocket conformation reproduced these critical interactions but was unable to appropriately score them, we sought to rectify this using the above induced fit method. To this end, CCR2 PDB 5t1a complexes docked with either **JNJ-27141491** or **CCX-140** were subjected to restrained conformational sampling with energy minimization to generate JNJ- and CCX-specific CCR2 allosteric pocket conformers. The resulting conformers predominantly differ from the parent structure PDB 5t1a in helical bundle expansion and sidechain rearrangement, including substantial reorientation of the solvent-facing K311^8.49^ and a lateral shift of TM7 (**Fig. 8B**). We find that this reorientation is necessary to accommodate larger chemotypes. Selected computationally generated models, called xCCX and yJNJ, had much improved recognition capacity via FA|FA ranking for their respective chemotypes. The model xCCX significantly improved recognition for CCX, GSK and BMS compounds, at the expense of JNJ and to a lesser extent AZ chemotypes (**Fig. 8C**). Conversely, yJNJ had improved JNJ and AZ recognition, at the expense of CCX, GSK, and UCB (**Fig. 8D**). Encouragingly, the ensemble of PDB 5t1a, xCCX, and yJNJ greatly improved FA|FA recognition of CCX, JNJ, and GSK chemotypes (poorly recognized by 5t1a alone) without a loss of recognition towards UCB and BMS (**Fig. 8E**). The two models were also challenged with the original allosteric active-vs-decoy discrimination task. Both xCCX and yJNJ (**Fig. 8FG**) demonstrated improved FA|FA recognition of the original set of 50 diverse allosteric actives from decoys, compared to PDB 5t1a alone (**Fig. 7E**). Forming an ensemble with PDB 5t1a, xCCX, and yJNJ further enhanced recognition (**Fig. 8H**).

### Structural determinants of CCR2 allosteric antagonist affinity and selectivity

The predicted compound poses in the optimized CCR2 intracellular allosteric pocket conformers (**Supplemental Data 2**) delineate shared and unique structural determinants of CCR2 allosteric antagonist affinity. The “roof” and “walls” of the pocket are formed by aliphatic and aromatic residue sidechains (V63^1.53^, L67^1.57^, L81^2.43^, A241^6.33^, V244^6.36^, Y305^7.53^, F312^8.50^, Y315^8.53^) making it predominantly hydrophobic; the invariable halo- and/or methyl-substituted aromatic rings in all examined antagonists (**Fig. 9A-E**) interact with these residues(**Fig. 9F-K**), providing shape complementarity to the pocket. The favorable packing of bulkier sulfonamide antagonists (e.g. GSK series and **CCX-140**) against the roof of the pocket is enabled by the subtle helical bundle expansion in the computationally-generated xCCX and yJNJ conformers (**Fig. 8B**) and is comparable to the interaction by the allosteric inhibitor vercirnon bound to CCR9 (PDB 5lwe [136]). Closer to the opening, the pocket features three exposed backbone amide hydrogens on intracellular helix 8 (BW positions 8.48, 8.49, and 8.50). These amides hydrogen-bond with various acceptor groups of the compounds including the carboxylic acids in AZ and BMS series (**Fig. 9A-B, F-G**) (similar to the structurally characterized CCR2-RA-[*R*] [32]), the sulfonamide oxygens in the GSK series and **CCX-140** (**Fig. 9C-D,H-J**), or the critical 5-ester [126] in JNJ thioimidazoles (exemplified by **JNJ-27141491**, **Fig. 9E,J**). The exact pattern of hydrogen-bonding to 8.48-8.50 backbones varies between the compounds; for example, **JNJ-27141491** hydrogen-bonds to K311^8.49^ and E310^8.48^ distinctly omitting F312^8.50^(**Fig. 9J**). Additional compound acceptor groups (e.g. the extra carboxylic acid of the GSK compound **SD-24**, **Fig. 9D**) may hydrogen-bond the sidechain of R138^3.50^ on the opposite side of the pocket (**Fig. 9I**). The outward-facing sidechain of K311^8.49^ plays different roles with different antagonists: the smaller compounds **kettle04_33** and **brown16_38(-)** (**Fig. 9A-B**) are occluded by it from the outside (**Fig. 9F-G**); whereas the size of bulkier sulfonamide antagonists **CCX-140** and **SD-24** (**Fig. 9C-D**) necessitates a different conformation of K311^8.49^ and prevents their occlusion from solvent (**Fig. 9H-I**).

**Figure 9.**
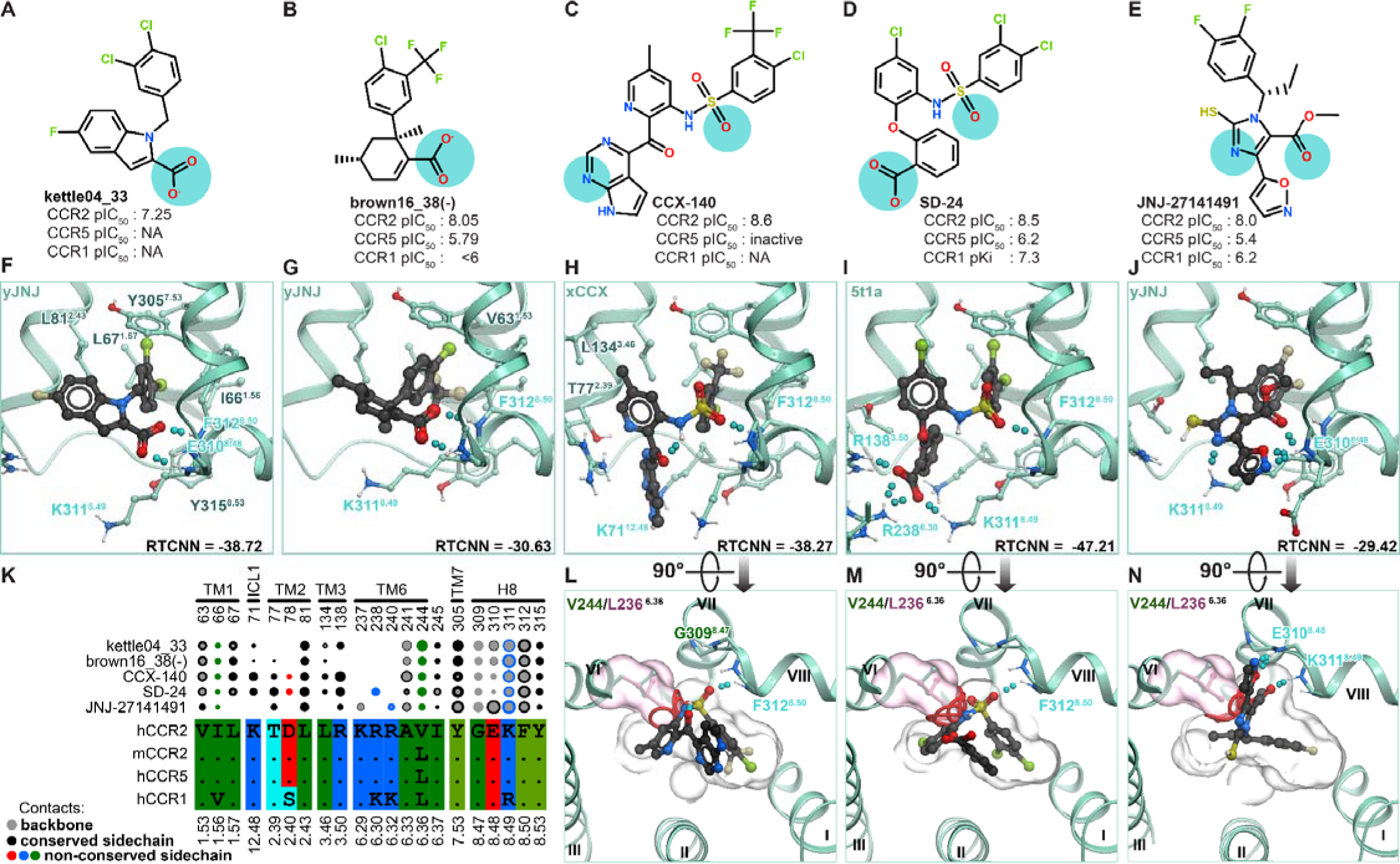
Antagonist pose analysis and structural determinants of allosteric affinity. (**A-E**) Chemical structures of clinically relevant CCR2 allosteric antagonists **kettle04_33** (**A**), BMS’s **brown16_38(-)** (**B**), **CCX-140** (**C**), GSK’s **SD-24** (**D**), and **JNJ-27141491** (**E**), and their experimentally measured potencies (expressed as pIC_50_) against human CCR2, CCR5, and CCR1. NA: not available. Cyan circles highlight atoms predicted to be involved in key polar interactions. (**F-J**) Predicted binding poses of compounds in (**A-E**) in human CCR2 allosteric pocket, viewed along the plane of the membrane. Key interacting amino-acid residues are depicted in sticks. Hydrogen bonds are shown as cyan dotted lines, receptor amino-acid residues involved in hydrogen bonding interaction are labeled in cyan. TM6 is hidden for clarity. (**K**) Partial sequence alignment and the map of the predicted contacts between compounds in (**A-E**) and amino-acid residues in the hCCR2 allosteric pocket. (**L-N**) Substituting V244^6.36^ for Leu makes the pocket sterically incompatible with the bound **CCX-140**, **SD-24**, and **JNJ-27141491**. Structures are viewed perpendicular to the plane of the membrane from the intracellular side. L244 (pink sticks and surface) collides with the Van der Waals radii of each compound atom as denoted by the red rings.

Hydrogen-bonding to the backbone amides in helix 8 appears universally critical for allosteric antagonist interactions with CCR2. The accessibility of these amides is imparted by the absence of the sidechain on CCR2 G309^8.47^ (**Fig. 9L-N**); these amides are also present in the homologous pockets of most other chemokine receptors because a glycine in BW position 8.47 is highly conserved in the chemokine receptor family (**Fig. S8**). This explains why CCR2 allosteric antagonists often have a weak but measurable secondary activity against CCR1, CCR5, or CXCR2 [121,123,137–139] but are completely inactive at non-chemokine GPCRs.

On the other hand, the exact shape of the hydrophobic “roof” and “walls” of the pocket appears to be important for allosteric antagonist selectivity within the chemokine receptor family. Mouse CCR2, as well as about a half of human chemokine receptors including CCR1 and CCR5, feature a V244^6.36^L substitution in the pocket (**Fig. 9K**, **Fig. S8**). Despite its seemingly conservative nature, introducing this substitution into the CCR2-antagonist complex models strongly alters the pocket shape and thus prevents favorable binding for all studied allosteric compounds (**Fig. 9L-N**). This is consistent with experimentally observed loss of activity of hCCR2 antagonists from the BMS, GSK, and JNJ series at human CCR5 and CCR1 [31,125,129,140]. Similarly, it explains why both **JNJ-27141491** [125] and **CCX-140** are almost inactive in mouse models despite their high potency in humans. In the case of GSK’s **SD-24**, the greater tolerance to the V244^6.36^L substitution - and hence stronger affinity for hCCR5 and hCCR1 (**Fig. 9D**) - may be attributable to the compensation by favorable hydrogen bonding with K71^ICL1^, R138^3.50^, R238^6.30^, and K311^8.49^ (**Fig. 9I**).

Altogether, the results of this section suggest shape complementarity and hydrogen bonding to helix 8 backbone amides as the key drivers of allosteric antagonist affinity to CCR2, and implicate a single conservative substitution, V244^6.36^L, as a determinant of hCCR2 antagonist selectivity against other human receptors and mouse CCR2.

## Discussion

A multitude of industry and academia drug discovery programs have yielded a broad array of potent CCR2 antagonists, and yet none have reached regulatory approval. This shortfall is attributed to various factors, including suboptimal affinities and residence time of compound candidates as well as species differences that complicate preclinical murine studies and the translation of results from animals to humans [29,47,48]. Compound discovery and property optimization can be rationalized and greatly accelerated by structure-based docking and screening, as demonstrated for numerous GPCRs and other target classes [141–150]. The prerequisites for such efforts include a good atomic-level understanding of the molecular determinants of affinity and selectivity of existing antagonists. Here we capitalized on the published experimental structures of antagonist- and agonist-bound CCR2 and CCR5 to develop such an understanding.

First and foremost, our results emphasize the importance of an accurate and predictive CCR2 binding pocket conformation for structure-based drug design. Despite the relatively subtle conformational differences between the orthosteric pockets of the active (chemokine-bound) and inactive (antagonist-bound) CCR2, only the pockets of inactive-state holo-structures were able to accurately dock antagonists or recognize them in virtual screening: in the active-state pockets, the recognition was effectively ablated. We attribute this to the collapse, in the active state, of the gap between receptor TM helices 1 and 7 that bridges the binding pocket with the surrounding lipid bilayer and, as our predictions show, is universally occupied by all known orthosteric CCR2 antagonists. Furthermore, we show that even between inactive-state pockets, VLS outcomes differ (consistent with the findings of others [151]) and are biased towards the ligand chemotype used for structure determination; binding poses of other chemotypes are recognized to a lesser extent, indicating the role of induced fit in antagonist binding. However, such biases may be surmounted by adapting the pockets to other chemotypes via conformational minimization. This process is a chemotype-specific implementation of an established practice called ligand-guided (or “ligand-steered”) structure refinement [105,152–159]. It enabled us to generate predictive models and derive accurate binding poses for orthosteric (BI), and allosteric (CCX, JNJ and GSK) compounds whose chemotypes are not captured by the available experimental structures. Importantly, a first-approximation pose obtained by docking into an inactive-state pocket was an essential precondition for such ligand-guided refinement.

We demonstrated that accurate pocket structures (like those from inactive antagonist-bound CCR2 structures or their optimized induced-fit derivatives) present exceptional opportunities for SBDD, due to their high predictive power in VLS. Indeed, despite a very small number of actives in the database (∼2% in the case of the master library, <1% for e.g. MK chemotype selective screening, etc.), all or almost all of them landed on top of the hit list, especially when ranked via an adequate scoring function. This high screening performance of experimental antagonist-compatible structures is in stark contrast with the reported performance of computational GPCR and chemokine receptor models, including those that were generated by the breakthrough AlphaFold2 technology [151,159,160]. The underwhelming predictive accuracy of AlphaFold2 models of GPCRs [151,160–162] may be because these models are found predominantly in active-like states and lack structural resolution in the binding pocket; the inability to predict induced fit and generate antagonist-compatible, high-accuracy, predictive pocket conformations may limit the drug discovery impact of AlphaFold2. However, if/when biased towards a conformational state appropriate to the scientific question, and able to predict even an approximately correct pose for even a single known compound, AlphaFold2 models can be further calibrated via ligand-guided refinement to make them usable for VLS and SBDD. Such refinement is not feasible when the first-approximation compound pose is inaccurate, as was the case with earlier homology models of CCR2 that, in spite of some successes in computational screening, in hindsight yielded poses inconceivable within the contemporary experimental structures [105,106,158,159]. Considering cryo-EM similarly struggles to resolve small molecule pocket conformation [163,164], ligand-guided structure refinement has enduring utility in producing VLS-suitable pockets from computationally and experimentally derived structures.

Next, we established the utility of an artificial intelligence-(AI-) based approach for antagonist docking and screening. AI-enabled scoring functions like RTCNN [83,84] or AtomNet [165] are highly effective at discriminating binders from non-binders in VLS applications. Compared to the traditional force-field-based full-atom score (FA), RTCNN excelled in our VLS assessments, especially when applied at the stage of library compound ranking (vs pose selection). In particular, it has readily overcome the problem of chemotype-selective recognition of CCR2 orthosteric antagonists by the two available experimental structures. By design, RTCNN has no conformational strain penalty and a weak steric collision penalty [166]; Consequently, it is more tolerant of small conformational differences within the binding pocket that otherwise lead to poor compound scoring by FA. This said, RTCNN was unable to sufficiently recognize the BI orthosteric chemotypes or the CCX and JNJ allosteric chemotypes in crystallographic pockets (as opposed to the computationally optimized induced fit pockets). Even more concerning, RTCNN could better “recognize” antagonists in a pocket whose geometry was completely inappropriate (e.g. the active-state G protein binding cavity for allosteric antagonists). Therefore the inclusion of physics-based measures remains critical. In our protocol, by using the physics-based FA scoring in compound pose selection, we ensured that conformational strain and steric interactions were respected prior to AI-based RTCNN scoring. It is important to note that both FA and RTCNN struggled to distinguish potent actives from less potent or inactive compounds of the same chemotype; this is expected as both of these scoring functions were developed for virtual screening and not for binding energy prediction. New AI-based binding energy prediction methods [167–171] or physics-based relative-free-energy methods [172–174] would be more appropriate for the binding affinity prediction task.

The identified (or generated) predictive CCR2 pocket conformations and the use of optimal docking and screening practices allowed us to reveal essential atomistic determinants of CCR2 orthosteric and allosteric antagonism. We demonstrated that orthosteric CCR2 antagonists universally rely on minor subpocket interactions including packing against the conserved W98^2.60^ and access to the inactive-state-dependent “tunnel” between helices TM1 and TM7; two out of three polar interaction determinants shared by all studied orthosteric CCR2 antagonists (Y49^1.39^ and T292^7.40^) are located in this tunnel. Our findings explain the previously reported detrimental effect of mutations to these residues [109,114,115] on compound affinity. We also identified the extra-helical, lipid-bilayer-facing groove accessible through this tunnel, and revealed its unexpected role for the binding of several antagonist chemotypes (e.g. Takeda and BI) not only to CCR2 but also to CCR5 (**Supplemental Data 1**), consistent with the shared conformational principles of activation and inhibition of these two receptors. Extra-helical targeting has been reported as a modulator mechanism for a number of GPCRs [175–180]; therefore, this previously uncharacterized CCR2 feature may potentially represent a new targetable interface for CCR2 (and, potentially, CCR5) antagonism. The obtained CCR2 antagonist binding poses account for experimentally observed affinity differences, highlight essential determinants of affinity through the per-atom RTCNN scoring, and thus enable structure-based SAR efforts.

Contrasting with CCR2’s orthosteric pocket, its intracellular allosteric pocket is considerably smaller, limited in the number and diversity of polar features, and completely unavailable in the active state of the receptor. The allosteric antagonists rely largely on shape complementarity and hydrogen bonding to the backbone amides of helix 8 for binding affinity. Consequently, there is significantly less latitude for compound potency or selectivity optimization. Through virtual screening, we revealed exceptional sensitivity of allosteric antagonists of various chemotypes to minute (chemotype-specific) changes in the pocket shape. Altogether, these findings provide a rationale for the challenges reported by various CCR2 drug discovery programs in improving the potency and selectivity of allosteric antagonists.

Last but not least, our work elucidates the mechanisms of CCR2 antagonist selectivity against mouse CCR2 and human CCR5. For the orthosteric case, through rational gain-of-function mutagenesis, we conferred CCR5 sensitivity to an otherwise CCR2-specific antagonist and thus demonstrated the critical role of the residue at BW position 2.63 in defining CCR2 vs CCR5 selectivity. Gain-of-function mutations, though difficult to realize, lend considerably more confidence than a comparable loss-of-function experiment. As the selectivity-determining minor pocket residue is conserved between human CCR5 and mouse CCR2, antagonists assessment in murine models is likely subject to the same selectivity mechanism, consistent with the wide-spread use of humanized mice for CCR2 antagonist assessment [89,118]. Our finding provides a way for rational design of compounds that either circumvent or exploit this mechanism to enhance selectivity for CCR2 or, vice versa, to develop dual-affinity antagonists. For the allosteric case, we identified a single amino-acid substitution (V-to-L in BW position 6.36) that is found in mouse CCR2 and many human chemokine receptors and that, despite its conservative nature, weakens or prevents the binding of allosteric hCCR2 antagonists to these receptors. These findings rationalize the published selectivity profiles of representative allosteric antagonists from different chemotypes and explain the difficulties that the ChemoCentryx labs experienced in trying to achieve high activity on mouse CCR2 in the allosteric series.

Overall, this work revealed the structural principles of CCR2 orthosteric and allosteric antagonism and produced receptor-ligand complex structures that can be used as starting points for future compound optimization. Importantly, we expanded the available CCR2 structural landscape by including computationally generated chemotype-specific receptor models that are well-suited for structure-based drug design.

## Methods

### Compound libraries

Compound structures and experimentally measured activities (mostly IC_50_ values) were retrieved from published literature, patents, and/or unpublished chemical databases at ChemoCentryx and BMS (**Supplemental Table 1**).

Master libraries for the active-versus-decoy assessment were generated manually emphasizing balanced representation of diverse chemotypes, CCR2 potency (pIC_50_>7), and pharmacological profiles. The master CCR2 orthosteric library consisted of 45 CCR2 antagonists of which 7 and 14 have been reported to have a comparable activity or be inactive against human CCR5, respectively. The master CCR5 orthosteric library consisted of 3 CCR5-selective and 7 dual-affinity antagonists. The master CCR2 allosteric library consisted of 50 diverse antagonists (pIC_50_>7); for most of these compounds, the activity against CCR5 or other chemokine receptors was unavailable. Compound diversity in the master libraries was assessed via UPGMA hierarchical clustering on Tanimoto distances of compound chemical fingerprints in the ICM molecular modeling software version 3.9-3b (Molsoft LLC, San Diego, CA). Property-matched decoys for either master library were retrieved as SMILES from the DUDe library [79] and converted to 2D in ICM.

Extended libraries of orthosteric and allosteric compounds used in chemotype-specific assessments were generated by incorporating expansive structure-activity datasets into the master libraries. The extended orthosteric library consisted of 330 BMS actives, 21 MK actives, and 83 BI actives of pIC_50_>7.5, plus the same inactives and decoys as in used in the actives-vs-decoys assessment. The extended allosteric library consists of 55 AZ, 10 BMS, 15 CCX, 49 GSK, and 30 JNJ actives of pIC_50_>7.0, 7 UCB actives of pIC_50_>6.5, and the allosteric actives and their decoys used in the actives-vs-decoys assessment.

Prior to docking and screening, all compounds were stripped of salts and hydrogen atoms, neutralized, and subsequently re-protonated at pH=7.0 using the pKa prediction model in ICM.

### Receptor structure preparation

Receptor structures were prepared using ICM 3.9-3b. Structures were obtained from the PDB [181]. Buffers, solvents, salts and detergents were removed; engineered stabilizing proteins fused in receptor ICL3 were deleted and so were Gαβγ and stabilizing Fab chains in active-state agonist-bound receptor structures. For CCR2b PDB 5t1a, engineered sequence 226-SRASKSRIPPSREKKA-240 was reverted to WT CCR2b 226-LKTLLRCRNEKKRHR-240; this included the restoration of WT identities to the allosteric pocket residues R237^6.29^ and K240^6.32^. For the CCR2a PDB structures 6pg[xs], four stabilizing mutations were identified and reverted to match WT CCR2b: C70^ICL1^Y, G175^4.60^N, and A241^6.33^D. For PDB 6gps, the 5th stabilizing mutation K311^8.49^E was reverted to its WT identity as well. The CCR5 stabilizing mutation G163^4.60^N found in PDB 7f1[qrst], 4mbs, 5uiw, and 6ak[xy] was also reverted to WT glycine. All newly introduced atoms in the mutated residues were assigned occupancy of 0. Structures were annotated with Ballesteros-Weinstein indices in ICM via the GPCRdb API and optimally superimposed by TM domain backbones. Ligand and histidine residue formal charges and protonation states were assigned via a pKa prediction model in ICM at pH 7. Structures were converted to the ICM format (internal coordinates) in the presence of the ligands, which included the addition of hydrogen atoms with subsequent optimization of orientations and geometries for polar rotatable hydrogens. In the process of conversion, the zero-occupancy atoms in aa residue side chains (including those introduced by restoring stabilizing mutations to WT identity) were subjected to a short Monte Carlo conformational optimization. Because of uncertainties in the conformation of K38^1.28^ in the CCR2 orthosteric pocket (side-chain disordered in PDB 5t1a), this residue in all CCR2 structures and its ortholog in all CCR5 structures were mutated to Ala.

### Virtual ligand screening

Screening was conducted using the VLS module in ICM 3.9-3b.

To determine the volume for orthosteric compound sampling, we used the superimposed experimental structure ensemble of CCR2 and CCR5 and calculated an envelope mesh that encompassed all small molecule antagonists within 5L and all chemokine residues within 10L from the BW residues 3.32 and 7.39 (the “floor” of the orthosteric pocket) in the respective receptors. For allosteric compound sampling, the envelope mesh was constructed around the crystallized pose of **CCR2-RA-[*R*]** (CCR2, PDB 5t1a [32]), preliminary docked poses of **kettle04_33**, **CCX-140**, **JNJ-27141491**, and **SD-24**, and crystallized allosteric compounds from other chemokine receptors: biaryl-sulfonamide **vercirnon** (CCR9, PDB 5lwe [136]), diamino-cyclobutene **00767013** (CXCR2, PDB 6lfl [182]), and thiadiazole-dioxide **cmp2105** (CCR7, PDB 6qzh [183]). Spaces for sampling of orthosteric and allosteric compounds were defined as 3D rectangular boxes encompassing the respective ligand envelopes with a margin of 4L in each direction.

Initial pose generation was done by docking library compounds into 3D grid-potential representation [81] of each receptor model calculated within an orthosteric or allosteric box defined as described above. For this, each library compound was converted to 3D and sampled in internal coordinates *in vacuo* to generate an initial set of diverse energetically favorable conformations; these conformers were then placed in the receptor binding pocket in four principal axes-aligned orientation to generate the initial compound pose “stack” after which favorable poses were generated via a biased probability Monte Carlo optimization with reactive history [80]. Top ten poses obtained from grid-based docking were combined with the full-atom receptor model and rescored using Full-Atom (FA), Mean-Force (MF), and Radial and Topological Convolutional Neural Network (RTCNN) methods. The ICM FA score is a GBSA/MM-type scoring function incorporating solvation, electrostatic, and entropic terms to approximate free energy of binding, augmented with a directional hydrogen bonding term, and accounting for ligand strain [81]. The ICM MF score is the statistical potential of mean force developed using a published knowledge-based methodology [82]. The ICM RTCNN score [83,84] is informed by a graph convolutional neural network that includes layers for topological (chemical graph) and 3D radial convolutions; the network was trained to recognize experimental and native-like complex geometries among decoy predicted favorably-scoring geometries. The RTCNN score does not incorporate any physical energy terms [83,84].

One top-scoring predicted pose for each compound was selected by FA, MF, or RTCNN and chosen to represent the compound. The multiple compounds in their selected poses were compared to each other and rank-ordered according to an alternative scoring function (also selected from FA, MF, or RTCNN), producing nine possible scoring schemes (FA|FA, FA|RTCNN, etc.). Receiver Operating Characteristic (ROC) curves were constructed using the calc3Rocs macro in ICM and visualized using the R ggplot2 library [184]. For ROC curves using ensembles of structures, docked poses for a single compound were pooled together across multiple models when rescoring, and the candidate selected as the best scoring of all models.

Final compound poses were visualized and rendered in the ICM suite 3.9-3b.

### Induced fit conformational optimization

Induced-fit conformers of CCR2 were produced in ICM 3.9-3b by minimizing receptor-ligand complex geometries taken from the initial virtual screen. For this, a smaller fragmented full-atom version of the receptor was built by first selecting residues within 5.5L proximity of the ligand and then iteratively expanding the selection to ensure that fragment length is no less than 9 aa (for orthosteric compounds) or 18aa (for allosteric compounds), and that proper disulfide bonds are included where applicable. For orthosteric complexes, the resulting individual fragments represented, separately, TMs 1, 2, 3, 4, 5, 6, 7, and ECL2 (a residue in ECL1 was deleted to ensure that TM2 and TM3 are represented by separate fragments). For allosteric complexes, the fragments represented TM1-ICL1-TM2 (contiguous), TM3, TM6, and TM7-H8; TM4 and TM5 fragments, which are not proximal to the allosteric pocket, were omitted to accelerate computation. Atoms omitted from the fragmented full-atom receptor were reintroduced via grid potential maps [81] to retain their physicochemical contributions to conformation whilst accelerating subsequent sampling.

Monte Carlo conformational optimization involved the ligand and all side-chains of the fragmented receptor. In addition, for orthosteric complexes, conformational sampling involved translational, rotational, and backbone torsional variables (i.e. full flexibility) of TM1, TM2, and TM7 (TMs 3-6 and ECL2 were kept stationary). For allosteric complexes, the fragments representing TM1-ICL1-TM2, TM3 and TM4 were kept stationary while TM6 and TM7-H8 were allowed to move. Positional harmonic restraints (“tethers”) were applied to ends of sampled helices to prevent excessive movement and preserve the gross geometry of the pocket. In addition, for the orthosteric case, backbone carbons of residues at BW positions 2.53, 2.56, 2.60, and 2.63 were tethered in place to prevent excessive movement of the TM2 fragment. Finally, distance restraints (min-max distances of 1.8L-2.4L and penalty weight of 1.0) were set to preserve intra-receptor and receptor-ligand hydrogen bonds; for the orthosteric complexes, these restraints involved Y^1.39^, Y^3.32^, Q^7.36^, E^7.39^, T^7.40^ and T^7.44^, while for allosteric complexes, the restraints were to the backbone amides of K311^8.49^ and F312^8.50^.

For orthosteric complexes, prior to global optimization, the system was subjected to gradient minimization for 10,000 steps. For the allosteric case, fragments representing TM1-ICL1-TM2, TM6, TM7-H8, and all receptor side-chains were subjected to gradient minimization for 100,000 steps; the ligand was kept frozen. After that, a global biased probability Monte Carlo simulation involving the ligands and the “mobile” receptor fragments (specified as above) was initiated and performed for 10^6^ steps. System conformations emerging from this simulation were clustered to produce 6-10 non-redundant conformers which were then tested in VLS.

The orthosteric complex ensembles aBI1, aBI3, bBI2, and bBI4 (**Fig. 3**) were generated from pre-docked complexes of **ebel13_22** in either the “a” or “b” chains of PDB 6gpx [63]. For allosteric pockets optimization, pre-docked receptor-ligand complexes of compounds **CCX-140** or **JNJ-27141491** bound to PDB 5t1a [32] were used for generating the xCCX and yJNJ optimized conformers.

### Plasmids and cloning

The CCR5 in pcDNA3.1 Hygro(+) was created by PCR amplification of the coding region of N-terminally Flag-tagged CCR5 with subsequent subcloning of the product into pcDNA3.1 Hygro(+) under the CMV (human cytomegalovirus) promoter [185]. N-terminally Flag-tagged CCR2 in pcDNA3.1 was described in [186]. The CAMYEL biosensor in pcDNA was originally described in [187] and optimized for increased luminescence output by introducing two mutations (C124A/M185V) in the luciferase part in [188,189]. WT rat Gαi3 in pcDNA3.1 [189] was generously gifted by the Ghosh lab, UCSD. CCR5 mutants were generated in the pcDNA-Flag-CCR5 background using the Q5 site-directed mutagenesis kit and primers designed using the NEBaseChanger tool from New England Biolabs. Mutants were cloned and amplified in *E. coli* strain XL10 Gold.

### Chemokines

CCL5 was produced as previously described [190]. Briefly, it was cloned into a pET21-based vector with an N-terminal His8 tag followed by an enterokinase recognition site, and insolubly expressed in BL21(DE3)pLysS *E. coli*. CCL5-containing inclusion bodies were collected by centrifugation, solubilized in a denaturing buffer containing 6 M guanidinium chloride, and the His-tagged chemokine was purified by Ni-nitrilotriacetic acid (NTA) affinity chromatography. Purified CCL5 was refolded by first adding 4 mM DTT to the eluate at pH>7, and then exchanging the eluate into a non-denaturing buffer (50 mM Tris, pH 7.5, 500 mM L-arginine, 1 mM EDTA, 1mM GSSG) with subsequent overnight incubation at 4 °C. The refolded product was dialyzed into a buffer consisting of 50 mM MES pH 5.7, 50 mM NaCl, treated with enterokinase at 37 °C for at least 8 hours to remove N-terminal His tag, passed again over Ni-NTA resin to separate the unwanted cleavage products, and finally purified by reverse phase HPLC on a C18 column. The collected CCL5-containing fraction was frozen, lyophilized, and stored at −80 °C. CCL2 was similarly expressed in *E. coli* BL21(DE3)pLysS cells and purified from inclusion bodies as described in [191].

For BRET assays, the powders were resuspended in sterile distilled deionized water, the concentration of the solution was quantified by absorbance at 280 nm and normalized to 100 μM for CCL5 and to 1 mM for CCL2.

### cAMP BRET assays

cAMP binding assays were performed using the BRET-based cAMP biosensor V2-CAMYEL [188,189]. HeLa cells (original stocks provided by the Ghosh lab, UCSD, La Jolla CA [189] were seeded in 6-well plates at 400,000 cells per well in MEM+10% FBS and transfected ∼6 hours later with 1 μg/well V2-CAMYEL, 750 ng/well of the WT or mutant chemokine receptor, and 750 ng/well of WT rGαi3 using the Mirus TransIT-X2 transfection reagent according to the manufacturer specifications. ∼24 hours after transfection, cells were washed with PBS, lifted with trypsin-EDTA, and re-seeded in a white/clear bottom 96-well plate at 40,000 cells per well in phenol-red-free DMEM with 10% FBS. ∼24 hours after reseeding, media was replaced with BRET assay buffer (PBS pH 7.4, 0.1% glucose, 0.05% BSA) and white backing tape was fixed to the underside of the 96-well plate. RLuc substrate coelenterazine-h and phosphodiesterase inhibitor 3-isobutyl-1-methylxanthine (IBMX) were added to the wells to a final concentration of 15 μM and 100 μM, respectively, mixed by shaking the plate, and incubated at room temperature for 2-3 minutes, following which BRET1 signal was measured with the TECAN Spark multilabel plate reader for 5 minutes to establish the baseline. For this, repeated luminescence reads were taken at 430-485 nm and 505-590 nm, and BRET was calculated as the emission ratio at 505-590 nm to 430-485 nm.

Next, for agonist EC_50_ experiments (**Fig. S6**), cells transfected with WT CCR5, CCR5(Y89S), and CCR5(F109H/S180P) were treated with 0.005-600 nM CCL5 (final in-well concentration) and CCR2-expressing cells were treated with 0.002-200 nM CCL2, followed by BRET reads for 5 min. All wells were then treated with 25 μM Forskolin and read for 10 minutes. AUC of the post-forskolin 1/BRET1 measurements was calculated and normalized to 0%-100% where 0% corresponds to the average of 2-6 wells with no response (containing no chemokine or the lowest concentration of chemokine) and 100% corresponds to 2-6 wells with the highest response (usually the highest concentration of chemokine) within the same experiment. Agonist concentration response curves were plotted and fitted with a two-parameter nonlinear regression (log[agonist] vs. normalized response with a variable Hill slope) in GraphPad Prism v.10.

For antagonist IC_50_ experiments (**Fig. 6**), after establishing the baseline BRET signal, cells transfected with WT CCR5, CCR5(Y89S), and CCR5(F109H/S180P) were treated with CCL5 at the final concentrations of 100 nM, 200 nM, and 10 nM, respectively, and CCR2-expressing cells were treated with CCL2 at 100 pM. This was followed by BRET reads for 5 min, the addition of 25 μM Forskolin as above, and another read for 10 minutes. Finally, wells were treated with a range of concentrations of CCR2-selective inhibitors **PF-4136309** (prepared from 20 mM stock solution in DMSO) and **cherney08a_22** (a.k.a. **BMS CCR2 22**, prepared from 100 mM stock in water), and BRET signal was read for 10 minutes. AUC of the post-antagonist 1/BRET1 measurements was calculated and normalized to 0%-100% where 0% corresponds to the average of 2-6 wells containing the highest concentrations of inhibitor and/or non-chemokine-stimulated samples and 100% corresponds to the average of 2-6 wells containing the lowest inhibitor concentration and/or antagonist-free samples. Antagonist curves were plotted and fitted with a nonlinear regression model (log[inhibitor] vs. normalized response with a fixed Hill slope of 1) using GraphPad Prism v.10.

## Author contributions

I.K. designed and supervised the research. J.R.D.D. and I.K. performed computational studies. G.M.W. cloned receptor mutants and performed cell-based pharmacological assays, under the supervision of I.K. P.Z., A.T., and P.H.C. generated and assessed the database chemicals. S.G. and T.S. generated receptor constructs; S.G., T.S., L.B.-R., and C.C.S. made chemokines, under the supervision of T.M.H.. J.R.D.D., G.M.W., and I.K. analyzed the data and wrote the paper. P.H.C., T.M.H., and P.Z. edited the paper. All authors approved the final version of the manuscript.

## Acknowledgements

We are grateful to Max Totrov, Eugene Raush, and Andrew Orry (Molsoft LLC, San Diego, CA) for the ICM software guidance. This work was supported by NIH grants R21 AI149369, R21 AI156662 (to I.K.) and R01 AI118985, R01 AI161880, R01 GM136202 (to I.K. and T.M.H.).

## Abbreviations

6gp[sx]: 6gpx and 6gps
6gpx[AB]: 6gpx chains A and B
6ak[xy]: 6akx and 6aky
7f1[qrst]: 7f1q, 7f1r, 7f1s, and 7f1t
AI: artificial intelligence
AUC: area under the curve
AZ: AstraZeneca
BI: Boehringer Ingelheim
BMS: Bristol Myers Squibb
CCR: CC chemokine receptor
CCL: CC chemokine ligand
FA: full-atom
CCX: ChemoCentryx
GPCR: G protein-coupled receptor
GSK: GlaxoSmithKline
hCCR2: human CCR2
[hm]CCR2: human or mouse CCR2
INCB: Incyte
JNJ: Johnson & Johnson
MF: mean force
mCCR2: mouse CCR2
ICM: internal coordinate mechanics
MK: Merck
PK: pharmacokinetics
ROC: receiver operating characteristic
RTCNN: radial and topological convolutional neural network
SBDD: structure-based drug discovery/design
TK: Takeda
TM: transmembrane
UCB: Union Chimique Belge
VLS: virtual ligand screening.

## Notes

### Competing Interest Statement

The authors have declared no competing interest.

### Summary of Updates

Sections on "Structural determinants of CCR2/CCR5 orthosteric antagonist selectivity" and "Structural determinants of CCR2 allosteric antagonist affinity and selectivity" revised to include new data; Figure 9 revised; Figures S1, S7, and S8 added, two authors added.

