## supplemental figures and tables for "Molecular determinants of antagonist interactions with chemokine receptors CCR2 and CCR5"

^4^(current affiliation) Blueprint Medicines, Cambridge, MA, USA

^5^(current affiliation) Lycia Therapeutics, South San Francisco, CA

^6^(current affiliation) Avidity Biosciences Inc., San Diego, CA

*Corresponding author: Irina Kufareva, Ph.D., Associate Adjunct Professor, Skaggs School of Pharmacy and Pharmaceutical Sciences, University of California San Diego, 9255 Pharmacy Lane, MC 0657, La Jolla, CA 92093. Phone: 858-822-4163.

### Supplemental Figures and Legends

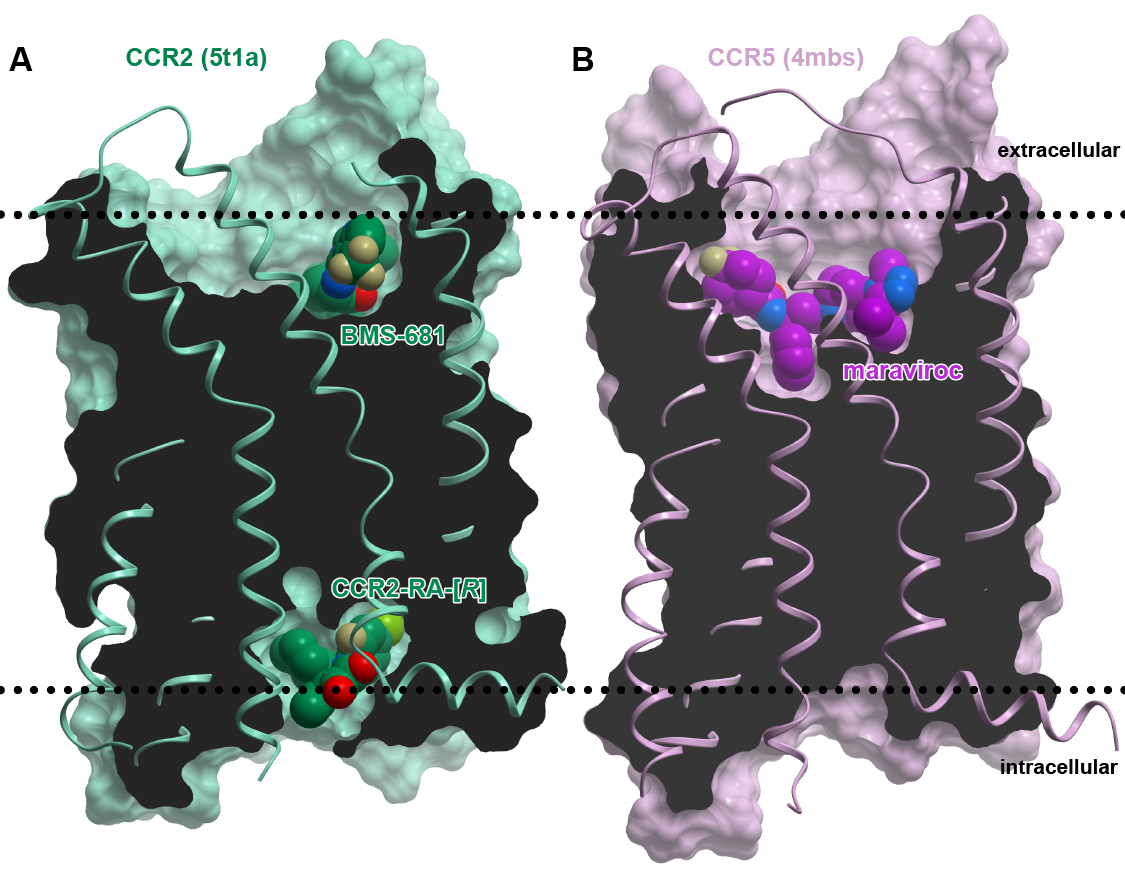

#### Figure S1. Experimental structures of CCR2 and CCR5 complexes with antagonists.

(**A**) CCR2 bound to **BMS-681** (which targets the extracellular orthosteric pocket) and the intracellularly targeted allosteric antagonist **CCR2-RA-[*R*]**.

(**B**) CCR5 bound to **maraviroc** which targets the extracellular orthosteric pocket.

Receptors are shown as ribbons and surface meshes and viewed parallel to the place of the membrane. The meshes are cross-sectioned to reveal the ligand binding cavities. Compounds are shown in a space-filling representation. Approximate membrane bilayer boundaries (extracellular and intracellular) are marked by dotted lines.

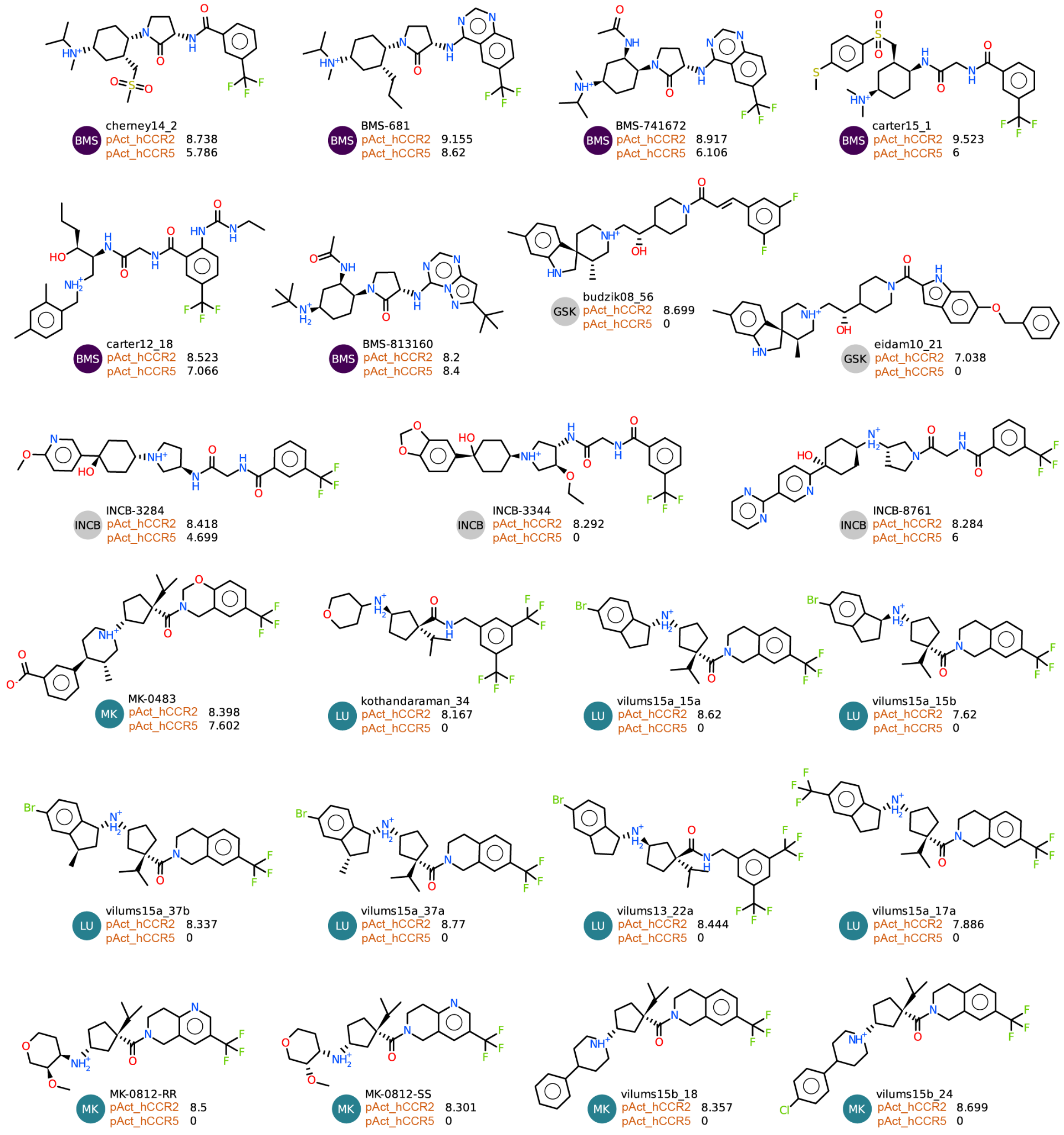

#### Figure **S**2**.** Structures of public-domain orthosteric CCR2 and CCR5 antagonists used in this study.

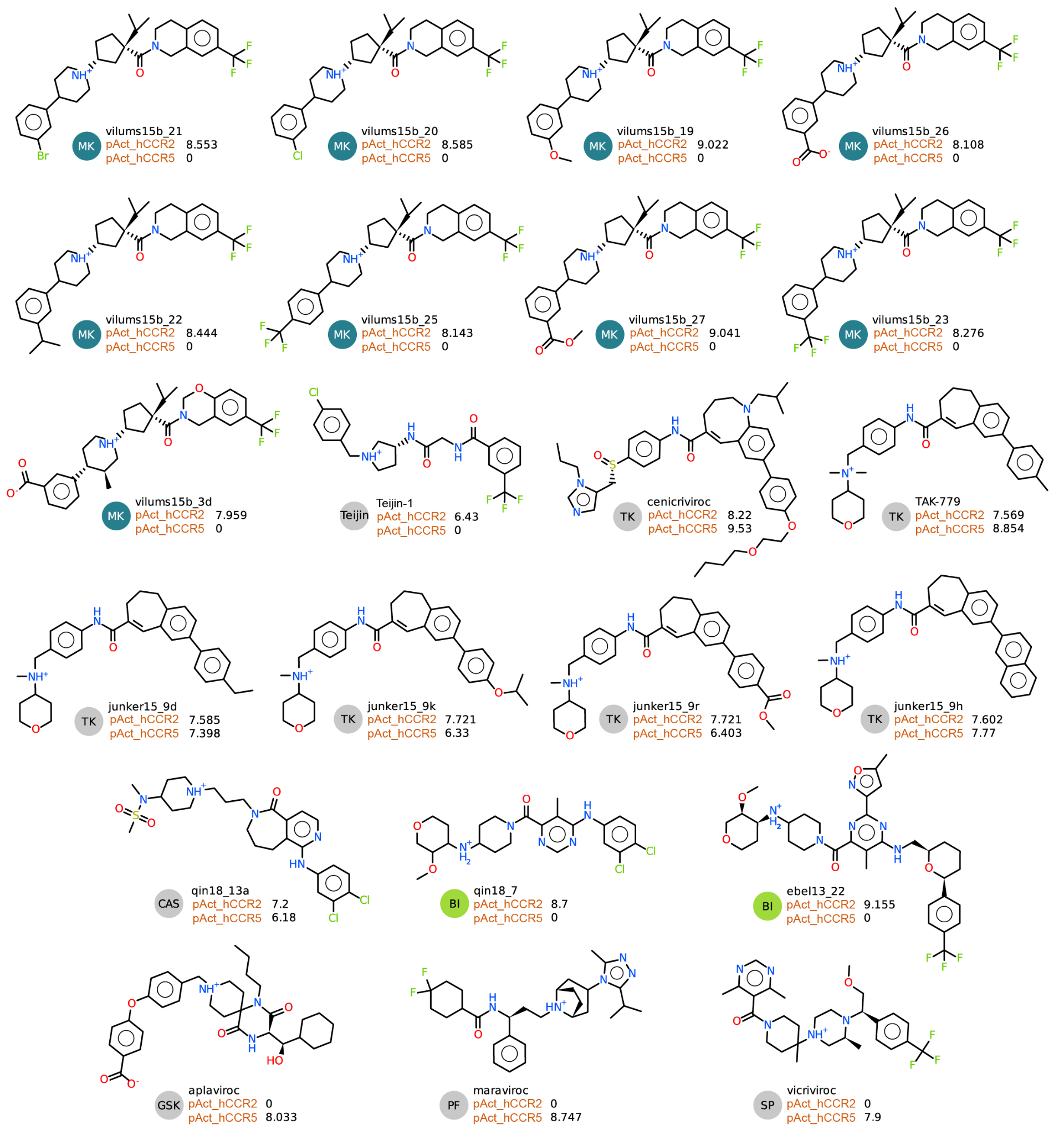

#### Figure S2 cntd. Structures of public-domain orthosteric CCR2 and CCR5 antagonists used in this study.

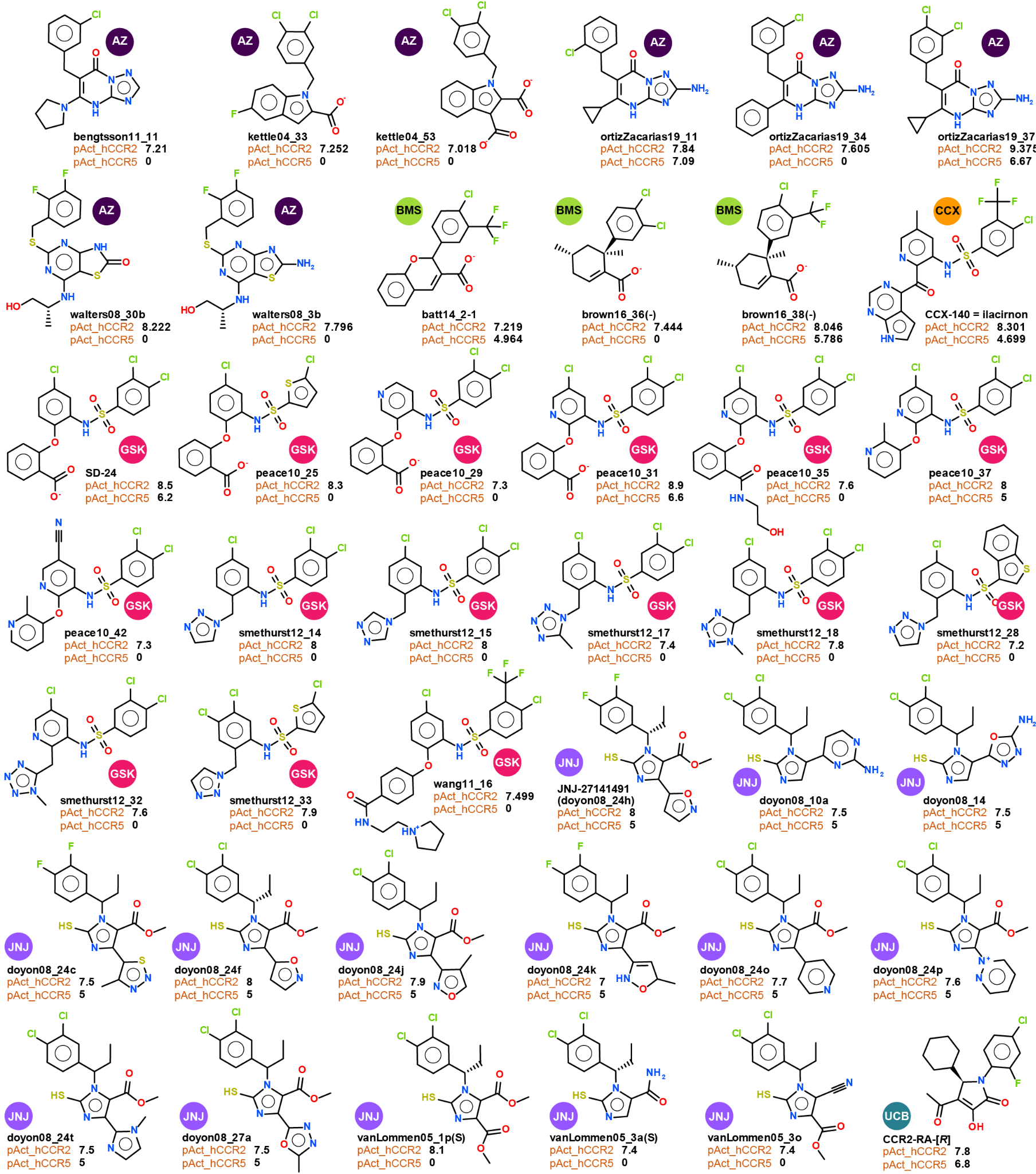

#### Figure **S**3**.** Structures of public-domain allosteric CCR2 antagonists used in this study.

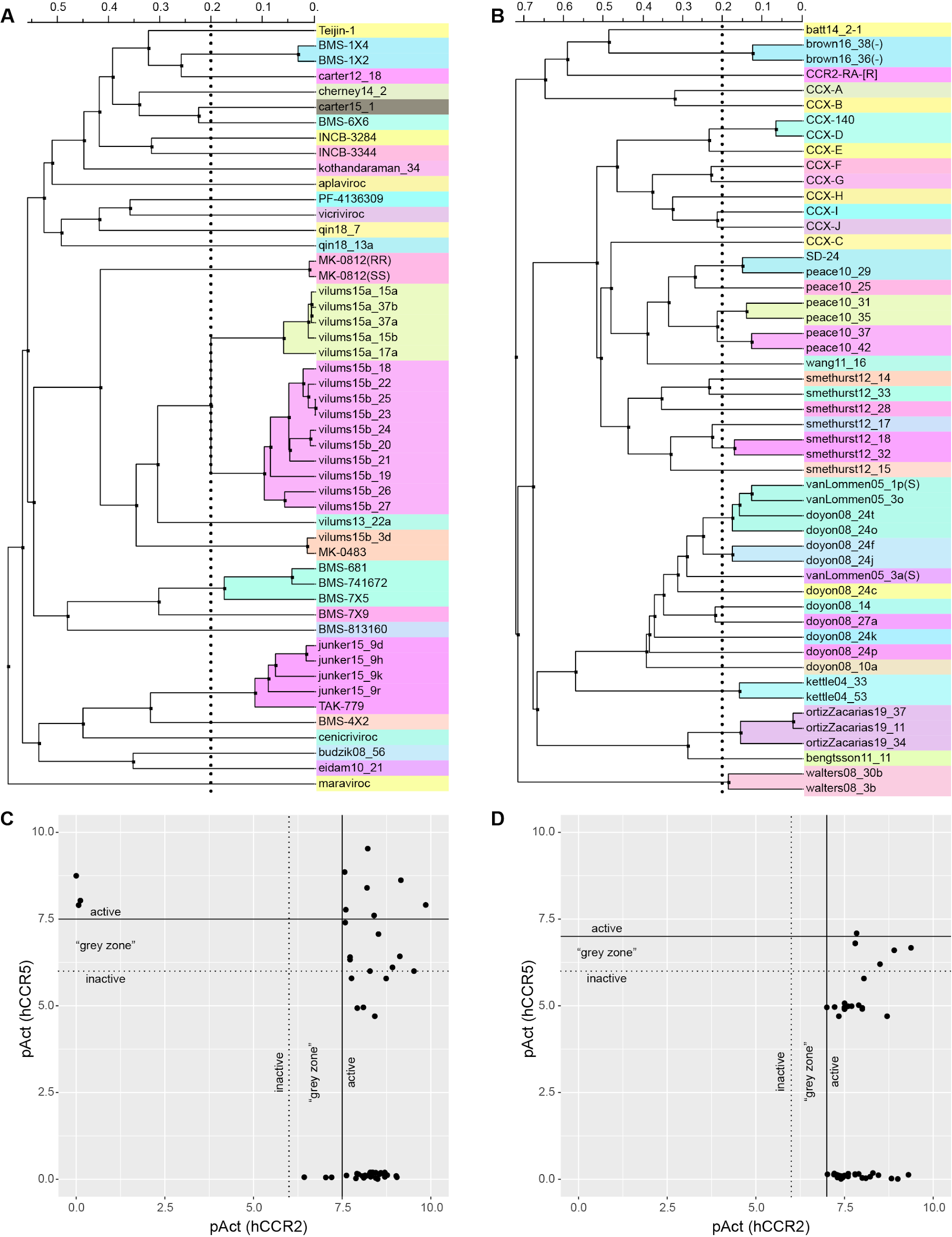

#### Figure **S**4**.** Compounds featured in this study are chemically and pharmacologically diverse.

(**A-B**) A hierarchical clustering dendrogram for the master library of 51 diverse orthosteric CCR2 and CCR5 antagonists (**A**) or the master library of 50 diverse allosteric antagonists (**B**) used in this study. Compounds were clustered by UPGMA based on Tanimoto distances of their chemical fingerprints as implemented in ICM. The dendrograms are colored by clusters emerging at the Tanimoto distance threshold of 0.2.

**(C-D**) Scatter plots of literature-derived inhibitory activities against human CCR2 vs human CCR5 for the 51 diverse orthosteric antagonists (**A**) or 50 allosteric antagonists (**B**) from the respective master libraries. Activities were expressed on a -log10 scale. Values close to zero indicate that the activity was not measurable or not available. Orthosteric antagonists with activities above 7.5 and allosteric compounds above 7 (solid lines) were considered active in all receiver operator characteristic (ROC) curves calculations. Orthosteric and allosteric compounds with activities below 6.5 or 6, respectively (dashed lines) were deemed inactive. Compounds within the “grey zone” between the solid and dashed lines were not regarded as active or inactive and were omitted when constructing ROC curves.

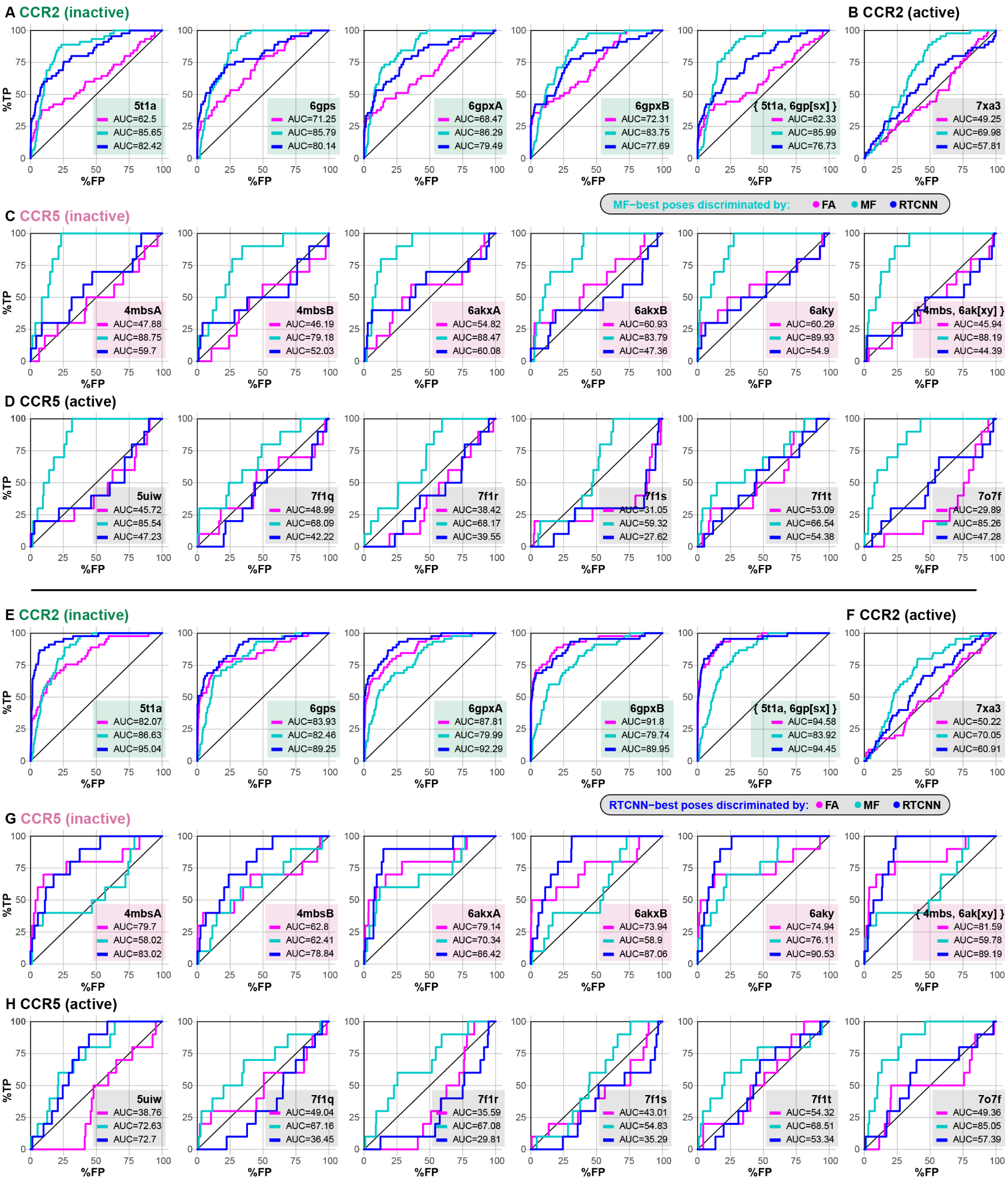

#### Figure **S**5**.** VLS recognition of known orthosteric antagonists by experimental structures of CCR2 and CCR5 when compound poses are selected using Mean-Force or RTCNN scoring.

(**A-B**) Receiver operator characteristic (ROC) curves obtained by VLS of a diverse library of 45 orthosteric CCR2 antagonists and a 54-fold larger set of property-matched DUDe decoys against the orthosteric pockets in inactive (antagonist-bound) (**A**) and active (chemokine agonist-bound) (**B**) CCR2 structure(s). For each docked compound, a single predicted pose was selected using the mean-force (MF) scoring function [[82]](https://paperpile.com/c/RwC4XM/bKwc) as implemented in ICM; compound ranking was performed using FA (magenta), MF (cyan), or RTCNN (blue).

(**C-D**) ROC curves obtained by VLS of a small diverse set 10 known CCR5 orthosteric antagonists and same inactives and decoys as in (**A-B**) against the orthosteric pockets in inactive (antagonist-bound) (**C**) and active (chemokine agonist-bound) (**D**) CCR5 structure(s). Pose selection and compound ranking were performed as in (**A-B**).

(**E-F**) ROC curves obtained by screening of the same library as in (**A-B**) against the same CCR2 structures, but with compound pose selection by RTCNN [[83,84]](https://paperpile.com/c/RwC4XM/v7He+xL7k).

(**G-H**) ROC curves obtained by screening of the same library as in (**C-D**) against the same CCR5 structures, but with compound pose selection by RTCNN.

#### Figure **S**6**.** Characterization of WT CCR2, WT CCR5, and mutant CCR5 responses to CCL2 and CCL5 in a cAMP suppression assay.

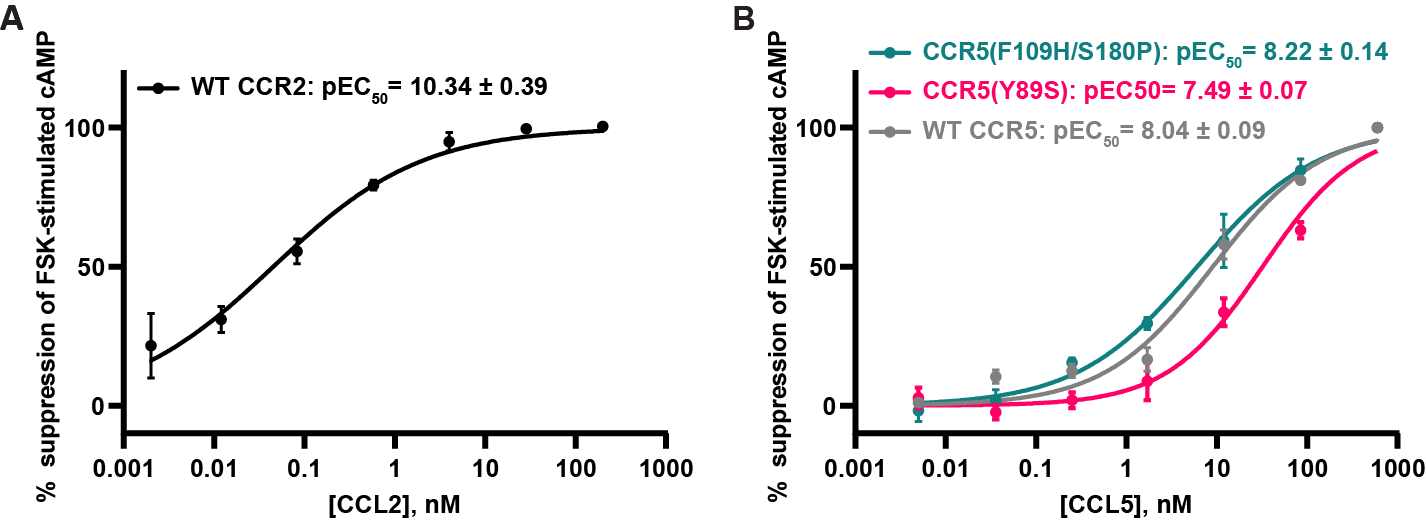

(**A-B**) Concentration-dependent suppression of forskolin-induced intracellular cAMP by CCL2 (**A**) and CCL5 (**B**) in HeLa cells transiently expressing WT CCR2 (**A**), WT CCR5 (**B**), CCR5(Y89S) (**B**), or CCR5(F109/S180P) (**B**). cAMP was measured via the BRET-based CAMYEL cAMP biosensor assay. Data within each experiment was normalized to 0%-100% as described in Methods. Error bars represent ±S.E.M. from n=3 independent experiments performed on different days, each with 2 technical replicates. Data was collated and fitted to a two-parameter nonlinear regression model (log[agonist] vs. normalized response with variable Hill slope) in GraphPad Prism.

#### Figure S7. Structural determinants of orthosteric selectivity in BMS acetamide-substituted cyclohexylamines [[118]](https://paperpile.com/c/RwC4XM/fSbZ).

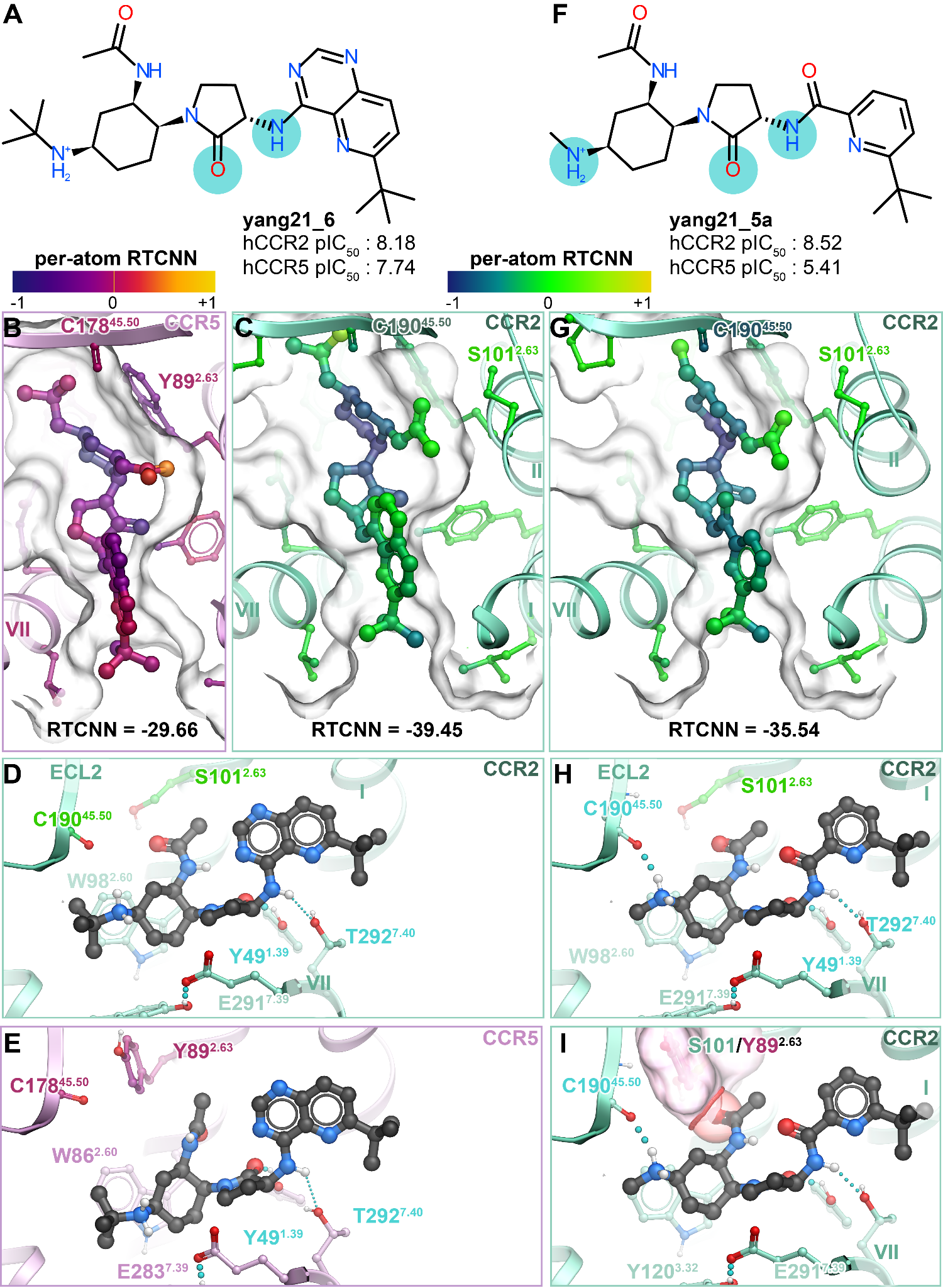

(**A-B**) Chemical structures and literature-derived receptor activities of dual-selective (hCCR2/hCCR5) BMS antagonist **yang21_6** and CCR2-selective antagonist **yang21_5a**. Cyan circles highlight atoms predicted to be involved in key hydrogen bonds.

(**C-E**) Poses of **yang21_6** in CCR5 (**C**) or CCR2 (**D**) and **yang21_5a** in CCR2 (**E**) are viewed perpendicularly to the plane of the membrane from the extracellular side. Atoms of the compounds are colored according to their contribution to RTCNN score.

(**F-I**) Predicted polar interaction networks for the compounds from (**A**-**B**) in the binding pocket of CCR2 (**F-G,I**) and CCR5 (**H**). Hydrogen bonds are shown as cyan dotted lines, receptor amino-acid residues involved in hydrogen bonding interaction are labeled in cyan.

(**I**) Steric hindrances highlighted between acetamide of **yang21_5a** and CCR5 Y89^2.63^ (pink surface).

#### Figure S8. Partial alignment of CCR2 allosteric pocket residues and its consensus across human Class A GPCRs and human chemokine receptors.

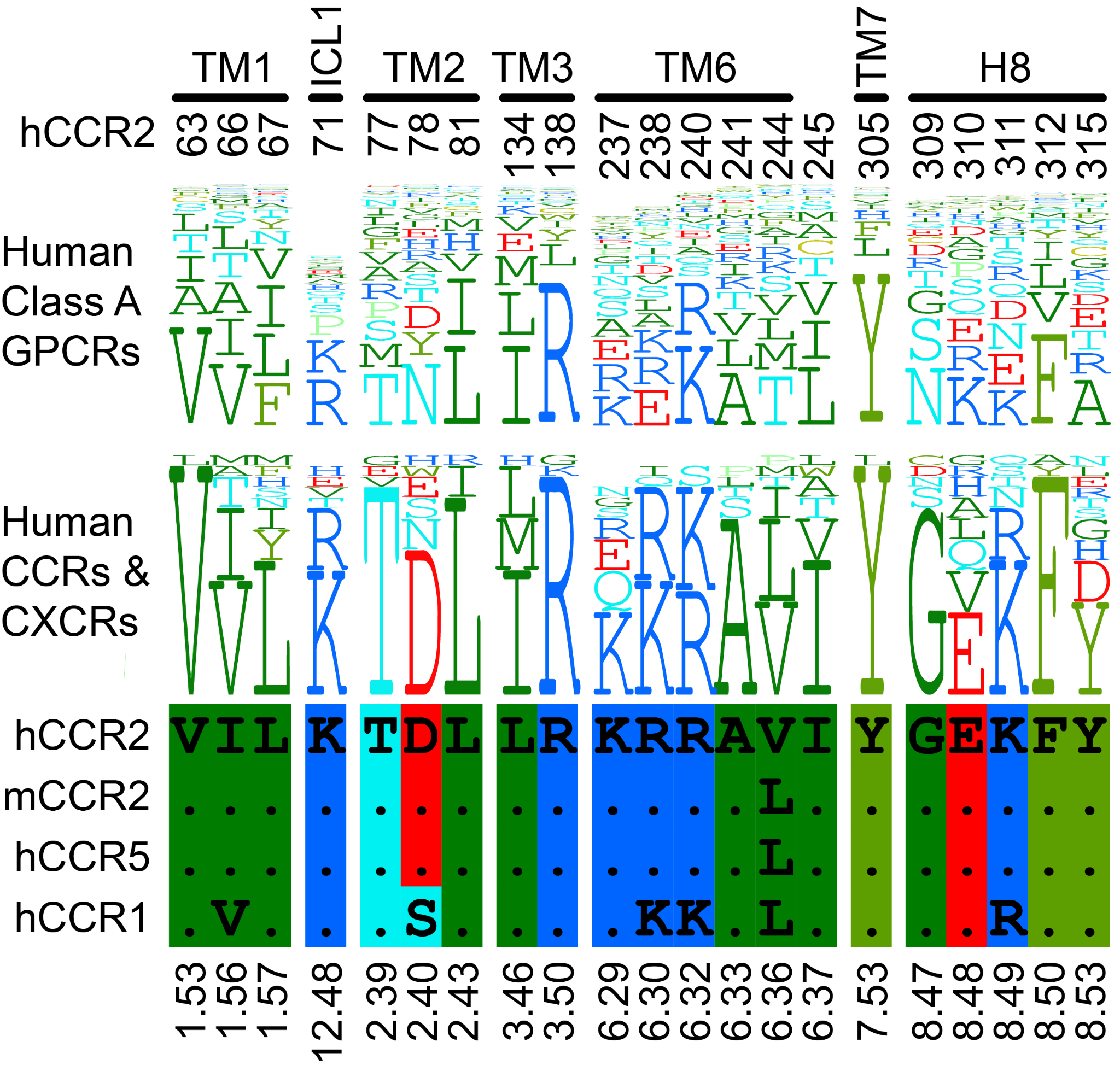

### Supplemental Tables

#### Supplemental Table 1. Overview of the chemical libraries of orthosteric and allosteric CCR2 antagonists used in this work.

| **Orthosteric** |  |  |  |  |
| --- | --- | --- | --- | --- |
| **Chemotype** | **# CCR2 / dual actives in the master library of 45* (pAct ≥ 7.5)** | **# cpds in the extended library** | **# CCR2 actives in the extended library (pAct cutoff)** | **References** |
| BI | 1 | 200 | 83 (7.5)  34 (8.65) | [[92,93,101]](https://paperpile.com/c/RwC4XM/29y3+i0Ow+Pu8M) |
| BMS | 12 / 3 | 530 | 330 (7.5)  65 (9.06) | [[73,87–91]](https://paperpile.com/c/RwC4XM/R6Zr+u8v9+T0r1+RyYC+E9yo+dBxv) |
| GSK | 1 | NA | NA | [[192]](https://paperpile.com/c/RwC4XM/FSCx) |
| INCB | 3 | NA | NA | [[68,69,102]](https://paperpile.com/c/RwC4XM/0XAH+k2b0+094z) |
| MK | 21 / 1 | 21 | 21 (7.5) | [[26,51,94–96,98,99]](https://paperpile.com/c/RwC4XM/xW5G+0b7S+iqVf+bafo+50H6+dMAz+oIeS) |
| TK | 7 / 3 | NA | NA | [[103,104]](https://paperpile.com/c/RwC4XM/XI9P+mgap) |
| **Allosteric** |  |  |  |  |
| **Chemotype** | **# CCR2 actives in the master library of 50 (pAct ≥ 7)** | **# cpds in the extended library** | **# actives in the extended library (pAct cutoff)** | **References** |
| AZ | 8 | 113 | 55 (7) | [[120–123]](https://paperpile.com/c/RwC4XM/kB9x+ztvU+A2GI+BCdm) |
| BMS | 4 | 50 | 10 (7) | [[129,130]](https://paperpile.com/c/RwC4XM/9iCu+t2Sd) |
| CCX | 9 | 40 | 15 (7) | [[131–133]](https://paperpile.com/c/RwC4XM/afnS+qhSD+mubV) |
| GSK | 15 | 79 | 49 (7) | [[128]](https://paperpile.com/c/RwC4XM/4nH3) |
| JNJ | 13 | 93 | 30 (7) | [[125–127]](https://paperpile.com/c/RwC4XM/iOoU+m45P+bOYd) |
| UCB | 1 | 52 | 7 (6.5) | [[119,124]](https://paperpile.com/c/RwC4XM/IIHG+nHUG) |

*Not listed in this table are 3 orthosteric antagonists with pAct(CCR2) <7.5 and 3 CCR5-selective orthosteric antagonists, for the total of 45+3+3=51 orthosteric antagonists in the master orthosteric library.

#### **Supplemental Table 2.** Reagents, supplies, and equipment used in the experimental part of this work.

| **Reagent** | **Source** | **Product Number** |
| --- | --- | --- |
| **Chemicals, Reagents and Recombinant Proteins** |  |  |
| PBS | Gibco | 20020023 |
| MEM | Gibco | 32561-037 |
| FBS | Avantor | 97068-085 |
| 0.25% Trypsin + EDTA | Gibco | 25200-072 |
| TransIT-X2 | Mirus Bio | MIR 6000 |
| BSA | Sigma-Aldrich | 9048-66-9 |
| Glucose | VWR | 97061-164 |
| Coelenterazine-h | RPI Corp | C61500 |
| IBMX | Sigma Aldrich | I5879 |
| Forskolin | Sigma Aldrich | F3917 |
| **PF-4136309** | MedChemExpress | HY-13245 |
| **cherney08a_22** a.k.a. **BMS CCR2 22** | Tocris | 3129 |
| DMSO | Sigma-Aldrich | D2650 |
| Q5 Site-Directed Mutagenesis Kit | New England Biolabs | E0554S |
| **Plasmids** |  |  |
| V2-CAMYEL | [[188,189]](https://paperpile.com/c/RwC4XM/LWHp+uG4W) |  |
| pcDNA3.1-Flag-hCCR2b | [[186]](https://paperpile.com/c/RwC4XM/FDa5) |  |
| pcDNA-Flag-hCCR5 | Cloned in-house [[185]](https://paperpile.com/c/RwC4XM/qypq) |  |
| pcDNA3.1-rGai3 | Ghosh lab, UCSD |  |
| **Primers (5’-3’)** |  |  |
| Y89S_fwd: TGGGCACATTCCGCAGCAGCC | Integrated DNA Technologies |  |
| Y89S_rev: GAATGGCACAGTGAGCAGAAAG | Integrated DNA Technologies |  |
| F109H_fwd: TGGCTTGTACAACATTGGCTTC | Integrated DNA Technologies |  |
| F109H_rev: GTCAGGAGCTGACACATG | Integrated DNA Technologies |  |
| S180P_fwd: TACTTGCTCCCCCCACTTCCCATAC | Integrated DNA Technologies |  |
| S180P_rev: TAATGCAGTCCCTCCTTC | Integrated DNA Technologies |  |
| **Other** |  |  |
| TC-treated 6-well plate | Falcon | [351146](https://ecatalog.corning.com/life-sciences/b2c/US/en/Cell-Culture/Cell-Culture-Vessels/Multiwell-Plates/Falcon%C2%AE-Plates/p/351146) |
| 96-well plate (white/clear bottom) | Greiner | 655098 |
| White backing tape | PerkinElmer | 6005199 |
| Spark Microplate Reader | TECAN |  |

#### **Supplemental** D**ata 1.** Orthosteric complexes featured in this work (3D coordinates in the PDB format)

#### Supplemental Data 2. Allosteric complexes featured in this work (3D coordinates in the PDB format)
